# Acyl-enzyme dynamics, tautomerisation and hydration regulate turnover of carbapenem antibiotics by the OXA-48 β-lactamase

**DOI:** 10.64898/2026.04.15.718664

**Authors:** Joseph F. Hoff, Michael Beer, Philip Hinchliffe, Catherine L. Tooke, Christopher J. Schofield, Marc W. van der Kamp, Adrian J. Mulholland, James Spencer

## Abstract

OXA-48 is a globally disseminated class D serine β-lactamase that efficiently confers resistance to a range of β-lactam antibiotics, including carbapenems, the most potent such agents versus *Enterobacterales* (*Escherichia coli* and relatives). Here we characterise the interactions of OXA-48 with the acyl-enzyme complex intermediates formed on its reaction with the carbapenems meropenem and ertapenem using X-ray crystallography and molecular dynamics (MD) simulations. X-ray crystal structures identify acyl-enzymes in both the Δ1-imine and Δ2-enamine pyrroline tautomeric forms. MD simulations show the epimeric Δ2 tautomers of meropenem and ertapenem to more frequently adopt binding poses competent for hydrolysis, i.e. with an appropriate orientation of the carbapenem 6α-hydroxyethyl group and positioning of the water molecule required for deacylation; the results indicate that the Δ2 tautomers are preferred for deacylation over the Δ1-tautomer. MD simulations based on the crystal structures show that, compared to OXA-48, acyl-enzyme complexes of OXA-519 (a natural OXA-48 variant with a single Val120Leu substitution adjacent to the catalytic general base) more frequently sampled conformations favouring hydrolysis, or formation of the alternative β-lactone deacylation product. MD simulations of complexes derived from quantum mechanics/molecular mechanics (QM/MM) simulations show the meropenem-derived β-lactone product is better retained in the OXA-48 active site than hydrolysed meropenem, consistent with reversible β-lactone formation. Overall, our results demonstrate how acyl-enzyme tautomerisation, dynamics and hydration collectively modulate degradation of 1β-methyl carbapenems by class D β-lactamases of the OXA-48 group, and how subtle changes in active site structure potentiate such effects in the OXA-519 variant.

## Introduction

β-Lactam antibiotics remain a core component of the antimicrobial arsenal used to treat bacterial infections^1^. Due to their widespread usage, various resistance mechanisms have emerged to limit their efficacy, of which the most commonly observed in Gram-negative bacteria involve production of β-lactamases, enzymes that hydrolytically inactivate β-lactams^2^. Serine β-lactamases (SBLs), comprising classes A, C and D of the Ambler (sequence-based) β-lactamase classification system, use a catalytic serine to mediate β-lactam hydrolysis^3^.

The OXA-48-like class D SBLs are most often found in *Enterobacterales,* an order of bacteria that includes pathogens responsible for antimicrobial-resistant infections globally^4,5^. The wide dissemination of OXA-48-like SBLs is of particular clinical concern due to their promiscuous catalytic profiles that permit inactivation of many β-lactam antibiotics. Members of this family (including OXA-48) confer resistance to carbapenems, one of the last lines of defence against multi-drug resistant infections **(Fig. 1)**^6–8^.

**Figure 1:**
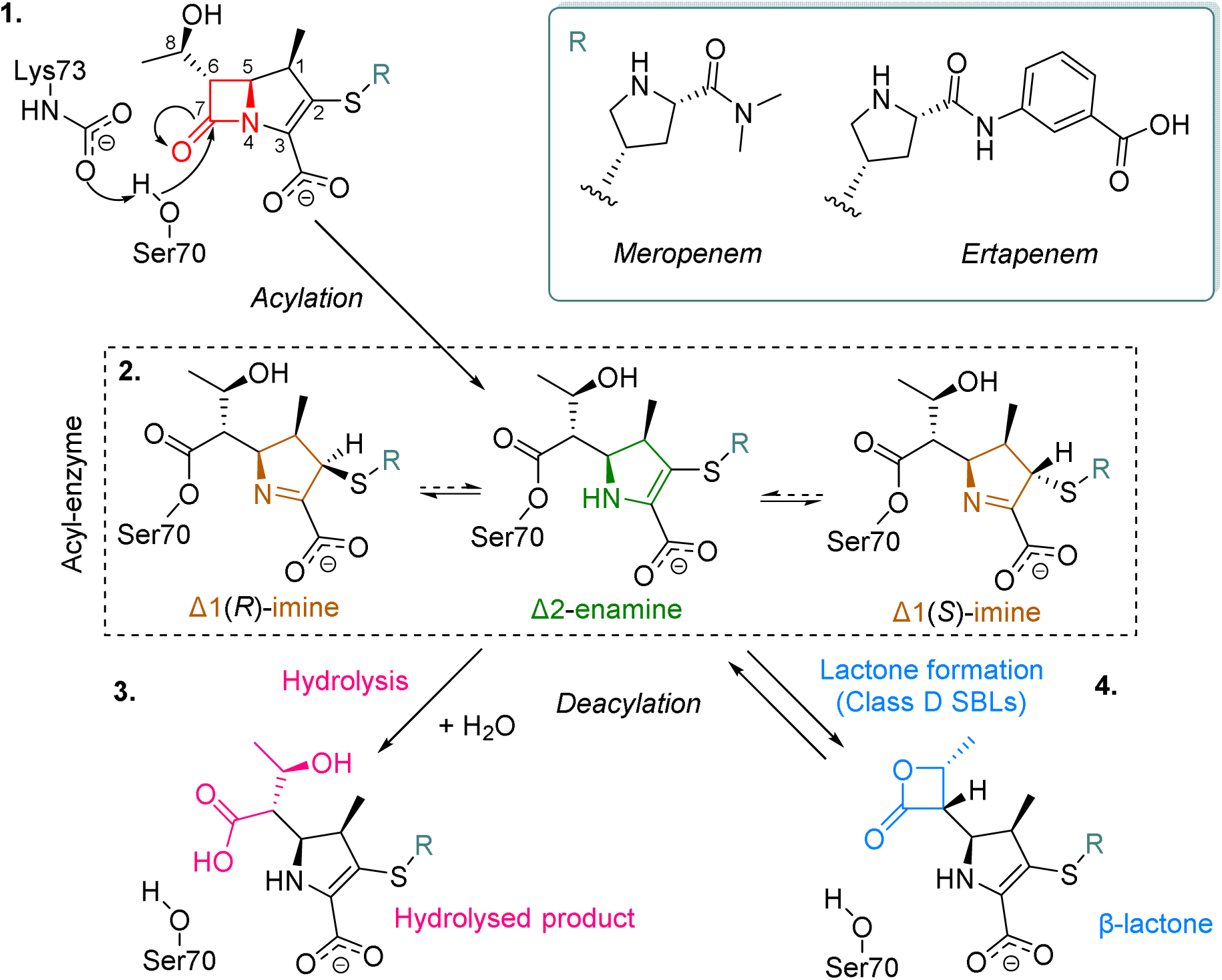
Breakdown of carbapenem antibiotics by class D SBLs. The β-lactam ring of a generic 1β-methyl carbapenem is shown in red, with active site residues numbered according to the OXA-48 sequence. C2 substituents (R-groups) of carbapenems studied here are shown in the adjacent box. During acylation (1), the activated nucleophilic Ser70 reacts with the β-lactam carbonyl carbon to form an acyl-enzyme complex. In the acyl-enzyme (2), the pyrroline ring can tautomerise between the Δ2-enamine (green) and the Δ1-imine (orange) in the R- and S-configurations. The carbapenem-derived acyl-enzyme breaks down to give hydrolysed or β-lactone products, through a water-mediated pathway (3) or via intramolecular rearrangement (4), respectively. For simplicity, only the Δ2-enamine tautomer is shown for the deacylation step.

SBLs hydrolyse β-lactams by a two-step acylation-deacylation mechanism **(Fig. 1)**. Class D SBLs are distinguished by use of a modified carboxylated lysine (KCX, Lys73 in OXA-48), adjacent to the catalytic serine, as the general base for both steps^2,9^. In acylation, the activated catalytic serine (Ser70 in OXA-48) attacks the electrophilic amide carbon of the β-lactam ring, resulting in a covalent intermediate, the acyl-enzyme. The acyl-enzyme is then resolved through a hydrolysis reaction mediated by a deacylating water molecule (DW) resulting in formation, and ultimately release, of a hydrolysed and inactivated product. Carbapenem-derived acyl-enzymes can tautomerise, via interconversion of the C2=C3 (Δ2-enamine) and N4=C3 (Δ1-imine) of the pyrroline ring isomers^10,11^. The Δ1-imine can also exist as one of two epimers, the Δ1(*S*) and Δ1(*R*) forms, which can interconvert *via* the tautomeric Δ2-enamine. In class A SBLs, the Δ2-enamine tautomer was identified as the catalytically competent form for carbapenem deacylation, which kinetic studies show to be the rate-limiting step in carbapenem breakdown by OXA-48-related enzymes^12,13^. NMR experiments suggest either the Δ2 or Δ1(*R*) forms of the carbapenem-derived acyl-enzyme to be favoured for deacylation in class D SBLs^11^. However, Δ2-enamine carbapenem-derived products rapidly tautomerise to the Δ1-imine forms in solution, making identification of the nascent product is challenging.

Class D SBLs can also deacylate carbapenem-derived acyl-enzymes to form β-lactone products, that are preferentially formed by carbapenems with a 1β-methyl substituent (e.g. meropenem, ertapenem)^14^. This occurs *via* intramolecular recyclization, without requirement for nucleophilic attack by a deacylating water. Here, the 6α-hydroxyethyl group of the carbapenem-derived acyl-enzyme is orientated for nucleophilic attack upon the C7 carbon, resulting in formation of the β-lactone ring. By contrast with hydrolysis, lactone formation is reversible, meaning that β-lactone product(s) in their respective tautomers can re-acylate and be subsequently hydrolysed^14^.

As in the hydrolytic reaction, the carboxylated Lys73 may act as a general base in β-lactone formation^14^. Aertker *et al.*^15^ demonstrated that changes in amino acid composition around the putative deacylating water channel, which houses KCX73, alters the amount of β-lactone formation in class D carbapenemases. One of these point mutations, Val120Leu, corresponds to a clinically observed OXA-48 variant, OXA-519, which was recovered from a multi-drug resistant *Klebsiella pneumoniae* isolate in 2015^16^. OXA-519 has a considerably modified catalytic profile, compared to OXA-48, with increased *K*_M_ values (suggesting lower affinities) for all classes of β-lactams, but a greatly enhanced turnover rate (*k*_cat_) for the 1β-methyl carbapenems meropenem (48.6-fold increase) and ertapenem (8.5-fold increase), which are turned over slowly by OXA-48 (**Table S1**)^16^. This corresponds to a decreased susceptibility to meropenem and ertapenem of *E. coli* TOP10 cells expressing OXA-519, compared to OXA-48. The enhanced activity of OXA-519 towards 1β-methyl carbapenems, compared to OXA-48, may be because it favours β-lactone formation as an alternative pathway for carbapenem degradation, although how the Val120Leu substitution in OXA-519 achieves this has been unclear.

Here, we report time-series of crystal structures of meropenem- and ertapenem-bound OXA-48 and OXA-519. Complexes with OXA-48 yielded both carbapenems as acyl-enzymes in both the Δ1 and Δ2 tautomers, with ertapenem in the previously unobserved Δ1(*R*) configuration. In contrast, OXA-519 acyl-enzymes were observed in the Δ2-enamine form only, at a range of time points, with Lys73 decarboxylated on extended incubation. We also observed meropenem and ertapenem hydrolysis products bound non-covalently to the OXA-519 active site. Extended molecular dynamics (MD) simulations (1.5 µs in total) of the OXA-48 ertapenem- and meropenem-derived acyl-enzymes, in all three tautomers, reveal the Δ2-enamine to be less mobile than other acyl-enzyme tautomers in the active site and to adopt a conformation that is more favourable for hydrolytic deacylation. This is evidenced by tautomer-specific orientation of the carbapenem 6α-hydroxyethyl group and therefore positioning of the deacylating water. Simulations of acyl-enzymes of OXA-519 showed that, compared to OXA-48, these can access more conformations suitable for deacylation by both the hydrolysis and β-lactone pathways. We attribute this to differences in the dynamics of the respective Val120 and Leu120 side chains, which affect hydration around the catalytic KCX73 base and the ability of the carbapenem 6α-hydroxyethyl group to sample conformations compatible with β-lactone formation. Control of the conformations, and interconversion between tautomers, of carbapenem-derived acyl-enzymes influences carbapenemase activity of OXA-48-like enzymes.

## Results

### Crystal structures of OXA-48 in complex with 1β-methyl carbapenems show multiple tautomeric states and 6α-hydroxyethyl conformations

Structures of OXA-48 (1.29 - 1.57 Å resolutions) in complex with meropenem and ertapenem were determined after soaking crystals for 1 hour, 2 hours and 16 hours (**Table S2A**). OXA-48 is a homodimer, and crystallises with two subunits (chains A and B) in the asymmetric (**Fig. 2A**)^17^. All structures presented here have similar backbone arrangements (RMSD < 0.31) when compared with a structure of a previously determined uncomplexed OXA-48, where crystals were grown in similar conditions (PDB 9H11^18^) (**Table S3**). OXA-48-bound meropenem and ertapenem were modelled as covalent acyl-enzymes, as evidenced by continuous *F*_o_-*F*_c_ electron density with Ser70 in the omit maps (**Figs. 2B, S1**). The presence of both the Δ2-enamine and Δ1-imine tautomers of both carbapenems could be identified by differences in the position of the C2-sulphur atom relative to the plane of the carbapenem pyrroline ring. In the 1 hour soak with ertapenem, the previously uncharacterised Δ1(*R*) configuration was observed in chain A, as indicated by strong electron density for the C2-sulphur atom in the same orientation, relative to the plane of the pyrroline ring, as the 1β-methyl substituent. In these structures, Lys73 is either carboxylated or it is decarboxylated and associated *via* its N^ε^-amine with either a chloride ion or water molecule (**Figs. 2B, S1 and Table S4A**). For the 1 hour structures, Lys73 is either fully carboxylated or in dual occupancy with the decarboxylated form. In contrast, after longer soaking times (16 hours for ertapenem, 2 hours for meropenem), Lys73 is fully decarboxylated. It should be noted that the pH of the crystallisation buffer differed between the 1 hour soaks and those at later time points (pH 8.8 vs 7.3 - 7.5, **Table S5**), and that this is likely to affect the carboxylation status of Lys73^19,20^.

**Figure 2:**
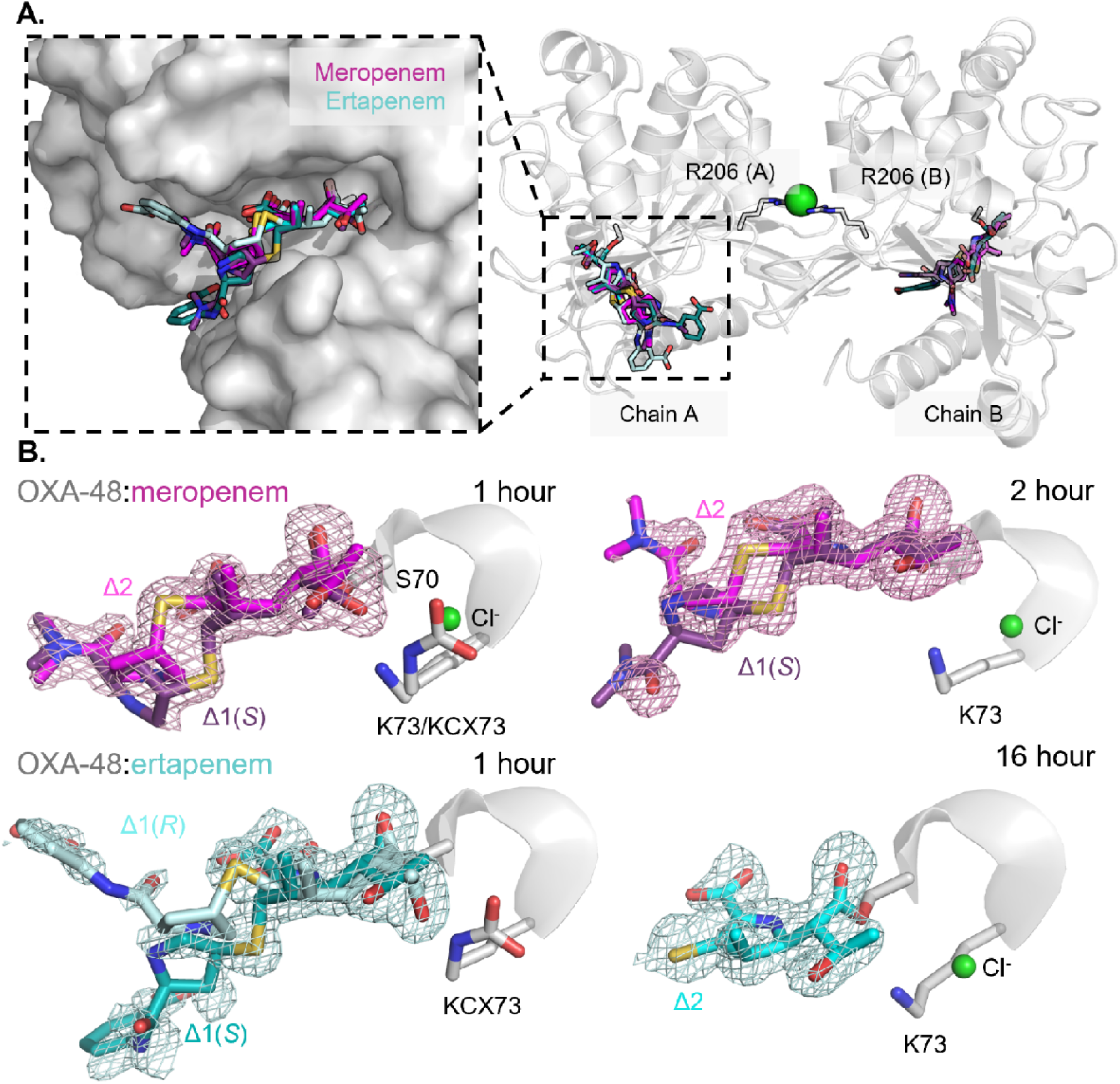
Crystal structures of meropenem- and ertapenem-derived OXA-48 acyl-enzyme complexes in Δ1 and Δ2 tautomers. (A) Structures of the OXA-48 homodimer (chain A light grey, chain B dark grey) bound to meropenem (pink/purple sticks) and ertapenem (cyan/blue sticks) as acyl-enzyme complexes. A chloride ion (green sphere) positioned at the dimer interface interacts with Arg206 from each chain (grey sticks). A magnified view of the active site (chain A) is shown in the dashed box (left) with overlaid acyl-enzymes shown as sticks. (B) F_o_-F_c_ omit maps (3σ, mesh) for meropenem and ertapenem (purple, cyan) acyl-enzymes (chain A, chain B sites are shown in **Fig. S1**). Modelled tautomers, assigned based on the position of the C2 sulphur atom (yellow), and Lys73 are indicated. For the data set collected after exposure to ertapenem for 16 hours, weak electron density prevented modelling of the C2 substituent beyond the C2 sulphur atom.

The core carbapenem ring derived atoms show binding poses that are similar across these complexes, as well as to those observed in previously published OXA-48 acyl-enzymes of ertapenem and meropenem^13,21,22^. In these complexes, the C7 ester carbonyl oxygen adopts an orientation pointing towards the oxyanion hole created by Tyr211 and Ser70, making hydrogen bonds with their backbone nitrogen atoms (**Fig. S2**). In addition, the C3 carboxylate group consistently forms a salt bridge with Arg250. In contrast, the C2 substituents of the meropenem- and ertapenem-derived acyl-enzymes show more variability in positioning, interacting with opposing faces of the OXA-48 active site (**Fig. 2A**). These interactions are summarised in **Note S1**. No modellable electron density was observed beyond the exocyclic sulphur atom, for the C2-subsituents of the Δ1(*S*) and Δ2 configurations of ertapenem-derived acyl-enzyme in the 16 hour structure.

In 3 of the 4 meropenem and ertapenem-derived acyl-enzyme structures, in chain B of the OXA-48 dimer there is a water molecule (W6) ‘wedged’ between the carbapenem C3 carboxylate and the pyrroline N4 nitrogen (**Fig. S3A-C**). W6 is also stabilised by hydrogen bonding networks with Thr209 and Arg250 and was only observed in carbapenem-derived acyl-enzymes with the Δ1(*S*) configuration. When overlaid with the active site of the opposing chain (chain A), in which W6 is not observed, its presence is associated with movement of the C3 carboxylate of ertapenem and meropenem, as indicated by an angle of rotation of 16.1 – 18.6° relative to the C6 carbon, which is static across the active sites (**Fig. S3D**).

Another point of variability in these structures relates to the orientation of the carbapenem-derived 6α-hydroxyethyl group. Previous work shows carbapenem-derived acyl-enzymes of class D β-lactamases to adopt one of the three dominant conformations of the 6α-hydroxyethyl group, as defined by their approximate C7-C6-C8-O dihedral angle: ∼50°, ∼180° and ∼290°, referred to here as conformations I, II and III respectively (using the nomenclature of Hirvonen *et al.*^23^). All structures of OXA-48:carbapenem acyl-enzymes (discounting complexes modelled as an inactive “flipped” acyl-enzyme intermediate^24^) currently in the PDB show this group modelled with dihedral angles ranging from 145° to 191°, i.e. grouped around conformation II (**Fig. 3, Table S6**). In conformation II, the oxygen of the 6α-hydroxyethyl group points away from the Lys73-carboxylate, out of hydrogen bonding distance. However, our structures of the Δ1(*S*) meropenem and Δ1(*R*) ertapenem complexes in the active sites of OXA-48 chain A at 1 hour soak times show the 6α-hydroxyethyl group to adopt an orientation closer to conformation III (∼290°), with C7-C6-C8-O dihedral angles of 321° and 279° respectively (**Fig. 2B**). In conformation III, Val120 is in an alternative side-chain rotamer (*g+*) to that seen in conformation II (*t*) and uncomplexed OXA-48 (PDB 9H11^18^), presumably avoiding steric clashes with the 6α-hydroxyethyl oxygen (**Fig. 3C**, panels 2 and 3)^15^. Conformation III of the 6α-hydroxyethyl group also corresponds with the presence of a carboxylated Lys73, with which the hydroxyethyl oxygen is in direct hydrogen-bonding distance (∼3 Å).

**Figure 3:**
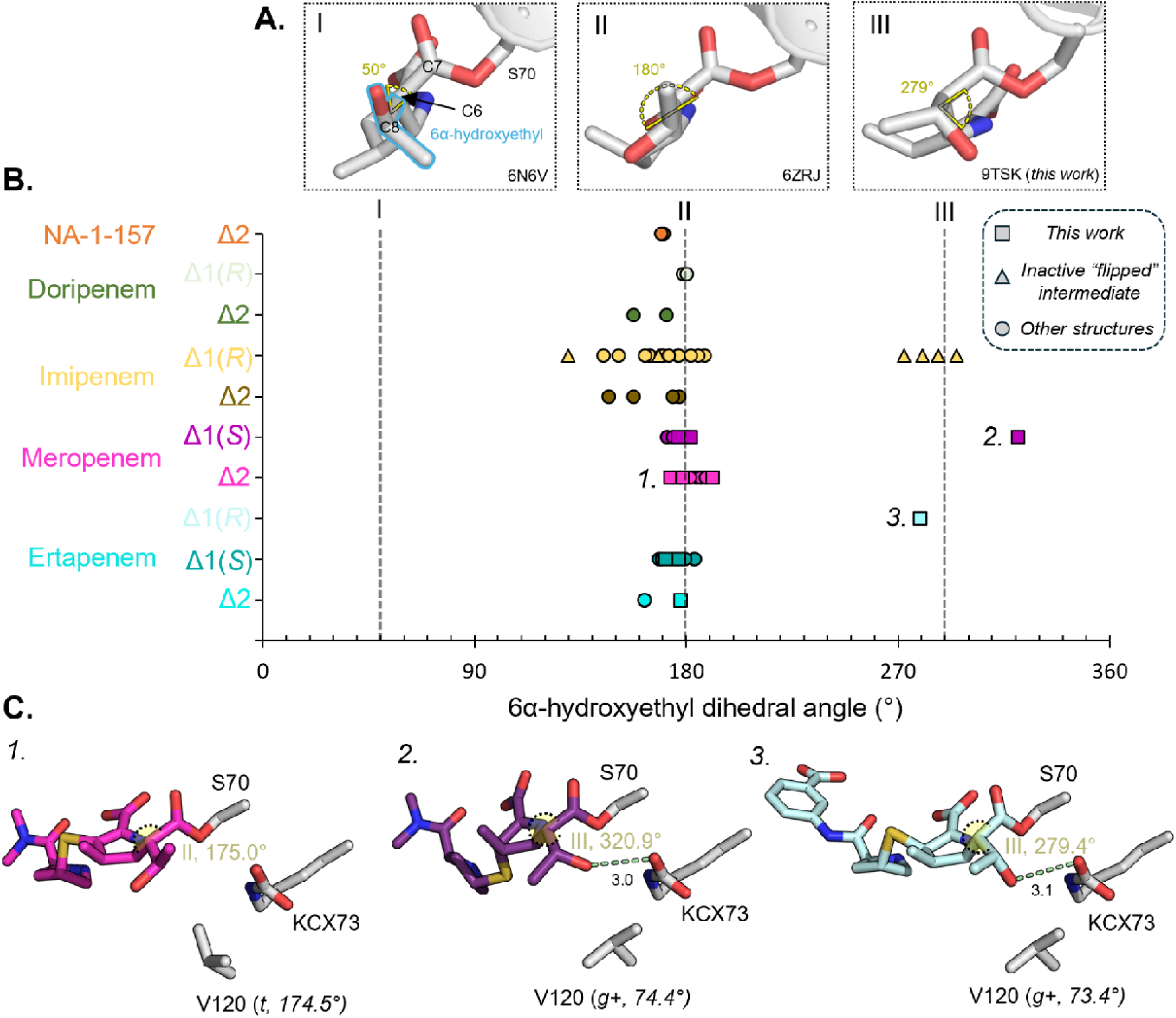
In crystallo conformational variation of the carbapenem 6α-hydroxyethyl group in OXA-48 acyl-enzyme complexes. (A) Representative 6α-hydroxyethyl conformations from crystal structures of class D SBLs in complex with carbapenem acyl-enzymes deposited in the PDB. Each 6α-hydroxyethyl dihedral (C7-C6-C8-O) angle is shown in yellow. (B) Distribution of the carbapenem C7-C6-C8-O dihedral angle (°) observed in all the OXA-48:carbapenem acyl-enzymes in this work and deposited PDB structures, with each angle represented as a dot (See **Table S6** for the full table of dihedral angles)^13,21,22,24–26^. The dots are separated for each carbapenem and tautomer, with those labelled 1., 2. and 3. corresponding to the structures in panel C. Triangles represent carbapenem-derived acyl-enzyme complexes in an inactive, “flipped” conformation where deacylation is prohibited (PDBs 7Q14, 7PFN)^24^. Squares are those introduced in this work and circles as other PDB-deposited structures. Grey dashed lines show 6α-hydroxyethyl conformations assigned by Hirvonen et al.^23^.(C) Active site views of OXA-48 (grey) bound to meropenem, modelled as: (1) Δ2 (magenta) and (2) Δ1(S)(purple) tautomer forms, and (3) ertapenem modelled in the Δ1(R) configuration (light blue), corresponding to the numbered dots in panel B. The hydrogen bond distance between the 6α-hydroxyethyl group oxygen of the reacted carbapenem and KCX73 carboxylate oxygen is shown as a light green dashed line. The Val120 side-chain rotameric form and the χ_1_ dihedral angle (N-Cα-Cβ-Cγ1) are in parentheses.

In contrast, no structures of the OXA-48:carbapenem-derived acyl-enzymes are available in which the 6α-hydroxyethyl is in conformation I (**Table S6**). This is consistent with the results of QM/MM simulations of the OXA-48 meropenem- and imipenem-derived acyl-enzymes, which show conformation I to be the most efficient orientation for hydrolytic deacylation^23^. It is therefore less likely for conformation I to be observed in crystal structures of carbapenem acyl-enzymes of class D carbapenemases. Although reported structures of other class D β-lactamases in complex with carbapenem-derived acyl-enzymes do show the 6α-hydroxyethyl group modelled in this orientation (**Fig. 3A**)^15^, these enzymes are either in a likely inactive form due to Lys73 being decarboxylated, e.g. OXA-23 carbapenemase bound to meropenem (PDB 6N6V^27^) and imipenem (PDB 6N6U^27^); or lack significant carbapenem-hydrolysing activity^28^, e.g. OXA-1 bound to doripenem (PDB 3ISG^29^).

### MD simulations of OXA-48:carbapenem acyl-enzymes show tautomer-specific control of deacylating water position and 6α-hydroxyethyl dynamics

Three independant 500 ns molecular dynamics (MD) simulations of the OXA-48:meropenem and ertapenem acyl-enzymes in each tautomer were run to investigate the dynamics of the different complexes (**Fig. S4A**). Per-residue root mean-squared fluctuation (RMSF) analysis shows no noticeable differences across the active site residues of OXA-48, with the biggest differences observed for the catalytic Ser70, to which the carbapenem-derived acyl-enzyme is covalently linked (**Fig. S5**). The (Ser70-bound) Δ1(*R*) tautomer of both the ertapenem- and meropenem-derived complexes fluctuates the most across these simulations, as indicated by average RMSF values of 2.93 Å and 1.57 Å, respectively, across both chains of OXA-48 (**Fig. S6**). The Δ1(*S*) configurations of the meropenem and ertapenem-derived acyl-enzymes undergo less extensive fluctuations, while the Δ2 tautomers of both carbapenems show the lowest average RMSF values (0.79 Å for the meropenem- and 1.04 Å for the ertapenem-derived complexes in the Δ2 configuration). RMSF analysis of carbapenem:Ser70 fragments shows these differences in atomic fluctuations to be predominantly associated with the extended C2 “tail” substituents of meropenem and ertapenem. Hydrogen bonding analysis of the core carbapenem scaffolds of each tautomer shows the greatest differences in interactions between the C3 carboxylate and OXA-48 active site, where both Δ1-imine enantiomers hydrogen bond much less frequently to Arg250 (Nη2) and Thr209 (Oγ1) than do the respective Δ2 tautomers (**Fig. S7**, hydrogen bonds 1 and 3). Previous crystallographic and simulation studies of the class A carbapenemase KPC-2 also demonstrated tautomer-specific differences in hydrogen bonding networks involving the C3 carboxylate of carbapenem-derived acyl-enzymes^12^.

The 6α-hydroxyethyl groups of meropenem and ertapenem in the different tautomers exhibit markedly different conformational dynamics in simulation trajectories (**Fig. S8A**). In the Δ2 tautomers of both the meropenem and ertapenem-derived OXA-48 complexes, the 6α-hydroxyethyl group predominantly resides in conformation I (50° dihedral angle), whereas in simulations of the Δ1(*S*) and Δ1(*R*) configurations this conformation is less populated, and instead conformation II (180°) is more frequently sampled. In all cases conformation III is very rarely sampled, as it is likely to be disfavoured due to steric hindrance from the 1β-methyl groups of meropenem and ertapenem^23^.

To understand how the orientation of the 6α-hydroxyethyl substituent, and the carbapenem tautomer, relate to meropenem and ertapenem deacylation, the distance between the carbapenem-derived C7 acyl-enzyme carbonyl carbon and the closest water molecule was measured over the trajectories (**Fig. 4**). 3 Å was selected as a distance cutoff for the deacylating water (DW) to be deemed as positioned to allow nucleophilic attack. Previous work^23^ has shown that in simulations of productive deacylation reactions, such an approach by the deacylating water costs less than 1 kcal/mol, consistent with frequent sampling of such positions in our MD simulations. For both the meropenem- and ertapenem-derived complexes, the closest water molecule is positioned for deacylation for the greatest proportion of the simulation time when the Δ2 tautomers are modelled (28.3% and 26.7% for the Δ2 configurations of meropenem and ertapenem respectively, averaged across both chains of OXA-48 (**Fig. 4A**)). This is followed by the Δ1(*S*) form of meropenem (8.5%) and the Δ1(*R*) form of ertapenem (16.1%). Finally, the meropenem-derived Δ1(*R*) and ertapenem-derived Δ1(*S*) tautomers support a water molecule in the deacylating position for the shortest amount of simulation time (8.1% and 8.0% respectively). Survival probability analysis shows that retention times for the deacylating water are also generally reduced in simulations of the meropenem and ertapenem OXA-48 acyl-enzymes in the Δ1-imine, compared to the Δ2-enamine, configurations (**Fig. S9**).

**Figure 4:**
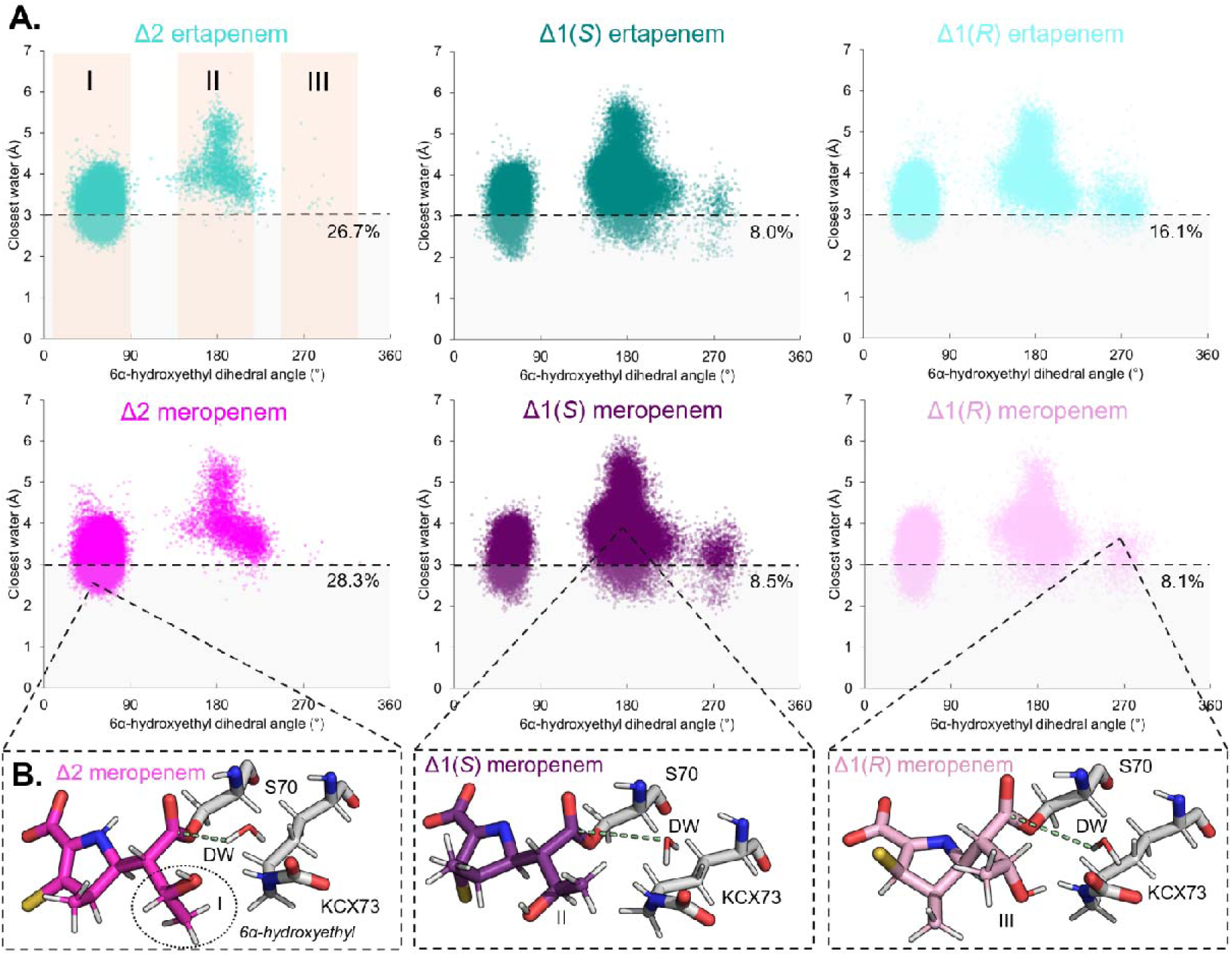
Relationship between carbapenem-derived acyl-enzyme tautomer configuration, 6α-hydroxyethyl conformation, and positioning of a deacylating water molecule, in MD simulations of carbapenem-derived complexes of OXA-48. (A) Scatter plots of carbapenem 6α-hydroxyethyl C7-C6-C8-O dihedral angle (°) against the distance of the closest water (any atom) to the carbapenem-derived acyl-enzyme C7 carbon over simulation trajectories of OXA-48 in complex with ertapenem (shades of blue) and meropenem (shades of pink) in Δ1(S), Δ1(R) and Δ2 tautomer forms. The dihedral angle spread for the 6α-hydroxyethyl conformations I, II and III are shaded in orange on the panel for OXA-48:ertapenem (Δ2). Each dot represents 40 ps snapshots over the 1.5 µs simulation time. The dashed line at 3 Å represents a distance cutoff for a water that would allow nucleophilic attack of the acyl-enzyme C7 carbon for deacylation. The percentage of simulations which have a closest water distance within 3 Å is shown beneath the distance cutoff line. (B) Representative simulation snapshots of meropenem-derived acyl-enzymes (pink sticks, C2-substituents beyond the sulphur are not shown) from the scatter plots above. KCX73 and Ser70 are shown as grey sticks, and the closest water to the acyl-enzyme carbonyl carbon highlighted as DW (for deacylating water). The carbapenem C7 distance to the closest DW atom is indicated by a pale green dashed line.

The conformation of the 6α-hydroxyethyl group in the meropenem and ertapenem-derived acyl-enzymes also appears to affect positioning of the deacylating water. In all cases, water molecules less than 3 Å away from the acyl-enzyme carbonyl carbon are most commonly observed when the 6α-hydroxyethyl group is in conformation I (**Table S7**). In general, conformation II shows a broader range of distances between the C7 carbonyl carbon and the closest water molecule, with fewer water molecules approaching to within the 3 Å distance deemed compatible with deacylation (**Fig. 4A**) suggesting that a potential deacylating water molecule is destabilised in this conformation. Indeed, inspection of simulation frames where the 6α-hydroxyethyl group is in conformation I shows the 6α-hydroxyethyl oxygen and the carboxylated Lys73 are both within hydrogen bonding distance of the deacylating water, therefore stabilising its positioning proximal to the acyl-enzyme C7 carbonyl carbon (**Fig. 4B**). In contrast, in conformation II, the hydroxyethyl oxygen would not be able to engage directly with the deacylating water, whilst the methyl group would likely sterically clash with a water molecule in this position. When the 6α-hydroxyethyl group is in conformation III, across all simulations, <1% of the water molecules closest to C7 are within the distance (3 Å) deemed compatible with deacylation (**Table S7**).

Class D carbapenem-derived acyl-enzymes are broken down through hydrolytic or β-lactone-producing pathways, representing competing routes for carbapenem degradation **(Fig. 1)**^14^. Therefore, we also considered the angle of nucleophilic attack (Bürgi-Dunitz trajectory^30^) for both carbapenem hydrolysis and β-lactone forming deacylation reactions, over the MD simulation trajectories. This was defined as the angle of the deacylating (i.e. closest) water molecule, or the carbapenem 6α-hydroxyethyl oxygen atom, relative to the acyl-enzyme C7 carbonyl for hydrolysis and β-lactone formation, respectively^14^. We note that the closest water molecule to the acyl-enzyme carbonyl C7 generally adopts an angle for nucleophilic attack that is further away from the optimum Bürgi-Dunitz trajectory (107°) than does the 6α-hydroxyethyl oxygen (**Fig. S10**). For both reactions, the Δ2-enamine forms of the carbapenem acyl-enzyme Bürgi-Dunitz angles are closer to 107° than is the case for the Δ1-imine forms^30^.

### β-Lactone carbapenem products are more easily retained in the active site of OXA-48 than the hydrolysis products

We next aimed to investigate the dynamics of different nascent active site complexed carbapenem-derived products, by modelling the meropenem-derived hydrolysis and β-lactone products, in their Δ2-enamine configurations, into one active site of the OXA-48 dimer using QM/MM simulations. Here, the C7 acyl-enzyme oxygen of both products was positioned within the oxyanion hole formed by the backbone nitrogen atoms of Ser70 and Tyr211 (**Fig. S11**). Three independent 500 ns MD simulations were then run under the same conditions as for the acyl-enzymes. In two of the three replicate simulations, the meropenem-derived hydrolysed product moves further away from Ser70 (**Fig. S12, Movie S1**). In the other replicate (Run 1), the hydrolysed product partially moves away from Ser70 before rebinding later in the simulation trajectory (**Movie S2**) with the C7 carboxylate returning into the oxyanion hole (**Fig. S12B**). In contrast, β-lactone-bound products were retained in the oxyanion hole across all simulations. In addition, they were less mobile in the OXA-48 active site, with hydrolysed meropenem having an average RMSF 0.48 Å greater than that of the equivalent β-lactone product across all three repeats. These results suggest that hydrolysed carbapenems are not retained in the OXA-48 active site as easily as are the carbapenem-derived β-lactone products, consistent with lactone formation by a reversible reaction^14^.

### Structural characterisation of OXA-519, an OXA-48 variant with enhanced carbapenemase activity, in complex with 1β-methyl carbapenems

We next aimed to structurally characterise an OXA-48 Val120Leu point mutant corresponding to the natural variant OXA-519, which has previously been shown to have enhanced hydrolytic activity towards meropenem and ertapenem and to more readily form β-lactone products than the parent OXA-48 enzyme^15,16^. Structures (1.43 – 1.95 Å resolution) of unliganded, meropenem- and ertapenem-bound OXA-519 were obtained from crystals grown under similar conditions to OXA-48 (**Fig. 5, Tables S2B-C**). OXA-519 has an almost identical backbone structure to OXA-48, and likewise assembles as a homodimer in the crystallographic asymmetric unit (**Table S3, Fig. S13A**). Both active sites of OXA-519 also have a very similar arrangement to OXA-48 (**Fig. S13B**). Lys73 is fully carboxylated in both structures (modelled with an occupancy of 1, **Table S4A**), and therefore the enzymes are likely to be in their active form under these conditions^9^. In both enzymes, residue-120 lines the so-called ‘deacylating water channel’ alongside Ser70 and Leu158, regulating solvent access to Lys73^21^. In previously deposited structures of uncomplexed OXA-48^17–19,21,24,31^, surface views indicate this channel to be completely closed due to the Cγ2 methyl group of Val120 (in the *t* rotamer) blocking access to the hydrophobic pocket where carboxylated Lys73 resides (**Fig. 5A, S13C**). In contrast, the deacylating water channel of OXA-519 is apparently much more open, and Leu120 does not occlude solvent access to Lys73 (**Fig. 5B**). Consequently, a water molecule (W2) is present deep within the active site pocket of OXA-519, but is not in OXA-48, likely due to steric clashes with Val120. This water is within hydrogen-bonding distance of the KCX73 carboxylate group (OQ1) and the side-chain oxygen of Ser70, therefore occupying a position that could facilitate water-mediated deacylation of β-lactam-derived acyl-enzymes.

**Figure 5:**
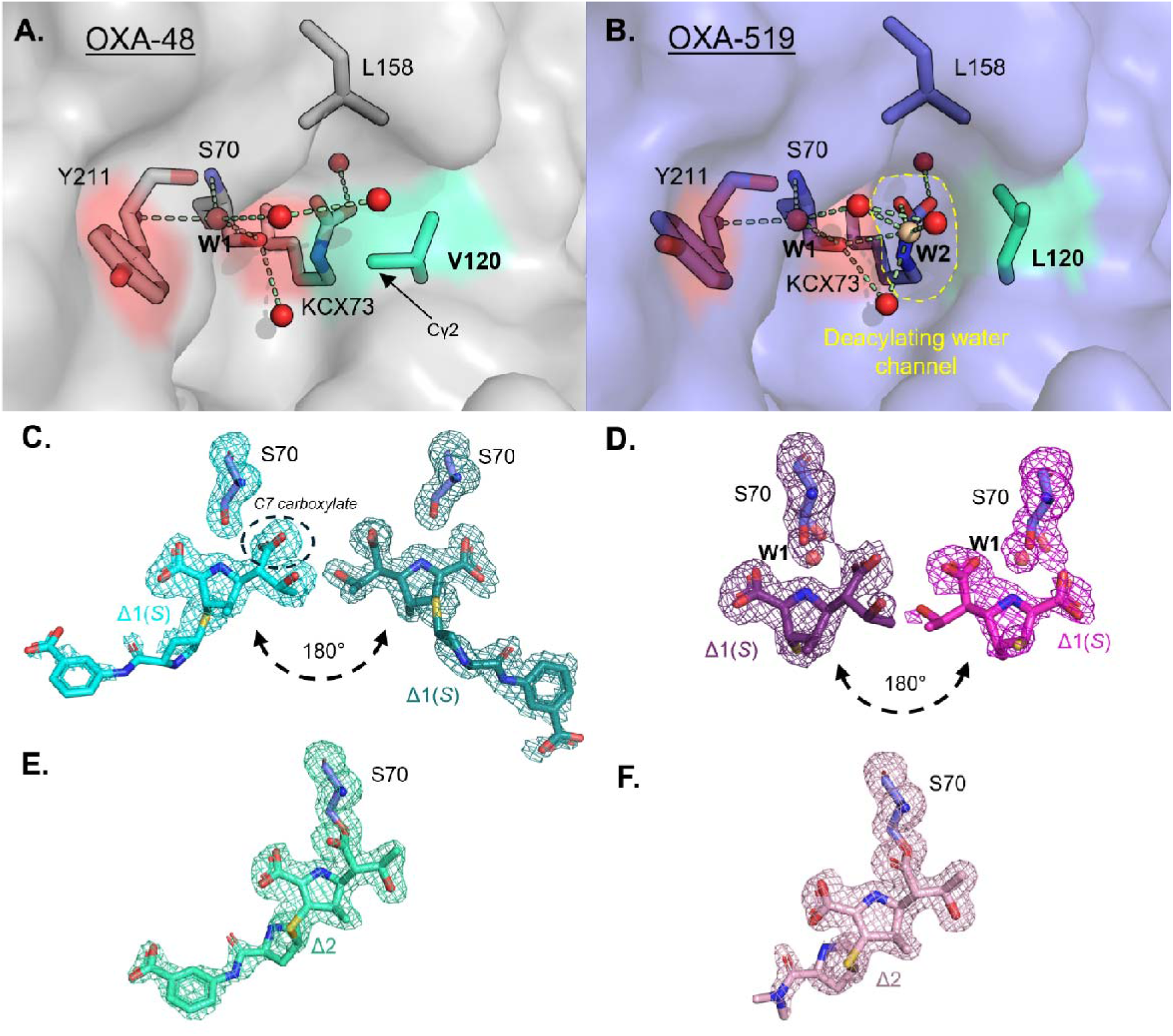
Crystal structures of uncomplexed and 1β-methyl carbapenem-bound OXA-519. Surface views of uncomplexed (A) OXA-48 (PDB 9H11^18^) (grey) and (B) OXA-519 (blue) deacylating water channels with waters conserved between these enzymes shown as red spheres (water in the oxyanion hole is labelled W1), and a water only observed in OXA-519 highlighted in beige (W2). Water-mediated hydrogen bonding networks are shown as green dashed lines. The outline of the open water channel in uncomplexed OXA-519 is highlighted by yellow dashed lines. Unbiased F_o_-F_c_ omit maps (mesh, contoured to 3σ) of (C) hydrolysed ertapenem (30 mins soak, (D) hydrolysed meropenem (2 hour soak), with different orientations taken from different active sites of OXA-519. (E) Ertapenem- (22 hour soak, chain A) and (F) meropenem-derived acyl-enzymes (22 hour soak, chain A) with OXA-519. Ser70 and dual-occupancy waters (W1, panel D) are included in omit selections. The pyrroline tautomer configurations of carbapenem-derived products and complexes are labelled.

We also determined structures of OXA-519 following incubation with the same concentrations of meropenem and ertapenem over a range of time-points (30 minutes, 2 hours, 4 hours and 22 hours; excepting the OXA-519:ertapenem structure after 22 hours incubation, for which a 10-fold lower concentration was used (**Tables S2B-C, S5**)). The resulting crystal structures show ertapenem and meropenem present either as the covalent acyl-enzyme intermediates, or as hydrolysed products bound to the active site pocket of OXA-519 in one of two orientations (**Fig. 5C-F**). In all cases, the hydrolysed products were modelled in the Δ1(*S*)-tautomer configuration, whereas ertapenem- and meropenem-derived acyl-enzymes are in the Δ2-enamine tautomer, with the C2-sulphur in the same plane as the carbapenem derived pyrroline ring. The meropenem and ertapenem-derived acyl-enzymes of OXA-519 show very similar binding poses and active site interactions to their equivalents formed with OXA-48 (**Fig. S14A-B**).

Hydrolysed products were modelled with occupancies ranging from 0.4 – 1.0, in dual occupancy with a water molecule that is present in unliganded OXA-519 (**Fig. S15 and Table S4B**). As noted above, the meropenem- and ertapenem-derived hydrolysis products are also observed bound in an orientation in which the molecule is “flipped” by ∼180° relative to the expected nascent hydrolysis product and the acyl-enzyme. Consequently, the C2 substituent of ertapenem-derived products (this region could not be modelled for meropenem-derived products) adopts strikingly different binding poses within the active site of OXA-519 (**Fig. S14C-F**). This binding pose is similar to that seen in reported crystal structures of OXA-10 (substituted with an OXA-48-derived β5 - β6 loop^32^) bound to hydrolysed doripenem and meropenem (PDB 6ZRG, 6ZW2). OXA-519 product complexes in both orientations likely represent re-bound products as the C7 carboxylate is positioned outside the oxyanion hole, with corresponding movement of Ser70, and are therefore not expected to represent the immediate end-points of enzyme-catalysed hydrolysis (**Fig. S16**). This contrasts with other reported structures of OXA-48 bound to hydrolysed imipenem (PDB 6PK0^22^, 7PEP^24^, 8QNZ^24^) in which the C7 carboxylate has remained within the oxyanion hole and the position of the Ser70 side-chain is unchanged.

In the structure determined at the earliest time-point (30 minutes), we observe a meropenem-derived acyl-enzyme in the Δ2-enamine configuration (**Fig. S17**). This complex also features a fully carboxylated Lys73 (**Table S4A**), suggesting that at this time-point OXA-519 is in a form that is active for carbapenem turnover, although the 6α-hydroxyethyl group is in a conformation (II, dihedral angle ∼180°)^9,23^ unlikely to support hydrolysis. At 2 hours, the hydrolysed meropenem products are observed (in both “flipped” and nascent orientations) with the Δ2 meropenem-derived acyl-enzyme re-emerging at 4 hours, at which point the Lys73 carboxylate group is present with partial occupancy (0.44 and 0.51 occupancies for chains A and B, respectively). For the ertapenem-derived complexes obtained at the same time-points, only the hydrolysed products are observed in the OXA-519 active site. Finally, for the 22-hour soaks, exposure to both carbapenems yields acyl-enzymes in exclusively the Δ2 configuration, with Lys73 fully decarboxylated.

### OXA-519:carbapenem acyl-enzymes sample deacylating conformations more frequently than OXA-48

MD simulations of the meropenem and ertapenem acyl-enzymes of OXA-519 in the Δ2-enamine configurations were subsequently run (**Fig. S4B**). Compared to the equivalent simulations of OXA-48, these show no major differences in RMSF (**Fig. S5**). We next filtered for simulation frames in which the acyl-enzyme adopts hydrolysis- or β-lactone-promoting (i.e. deacylation-competent) conformations, consistent with proposals in the existing literature^14,15,23^. These are characterised by limited hydration around KCX73^OQ^^2^ (one water molecule rather than two), with the carbapenem 6α-hydroxyethyl group in conformation I (**Fig. 6A**)^15,23^. For conformations supporting hydrolysis, a water molecule is additionally required to be proximal to KCX73^OQ^^1^ and the acyl-enzyme C7 carbon, whereas β-lactone-promoting conformations have the carbapenem-derived 6α-hydroxyethyl oxygen within hydrogen bonding distance (3 Å) of KCX73^OQ^^1^. In both cases, we also filtered for simulation frames in which the nucleophile (6α-hydroxyethyl oxygen for β-lactone formation or closest water oxygen for hydrolysis) is oriented with a Bürgi-Dunitz angle close to the optimal value (107±20°) for addition to the carbapenem C7 carbonyl^14,30^.

**Figure 6:**
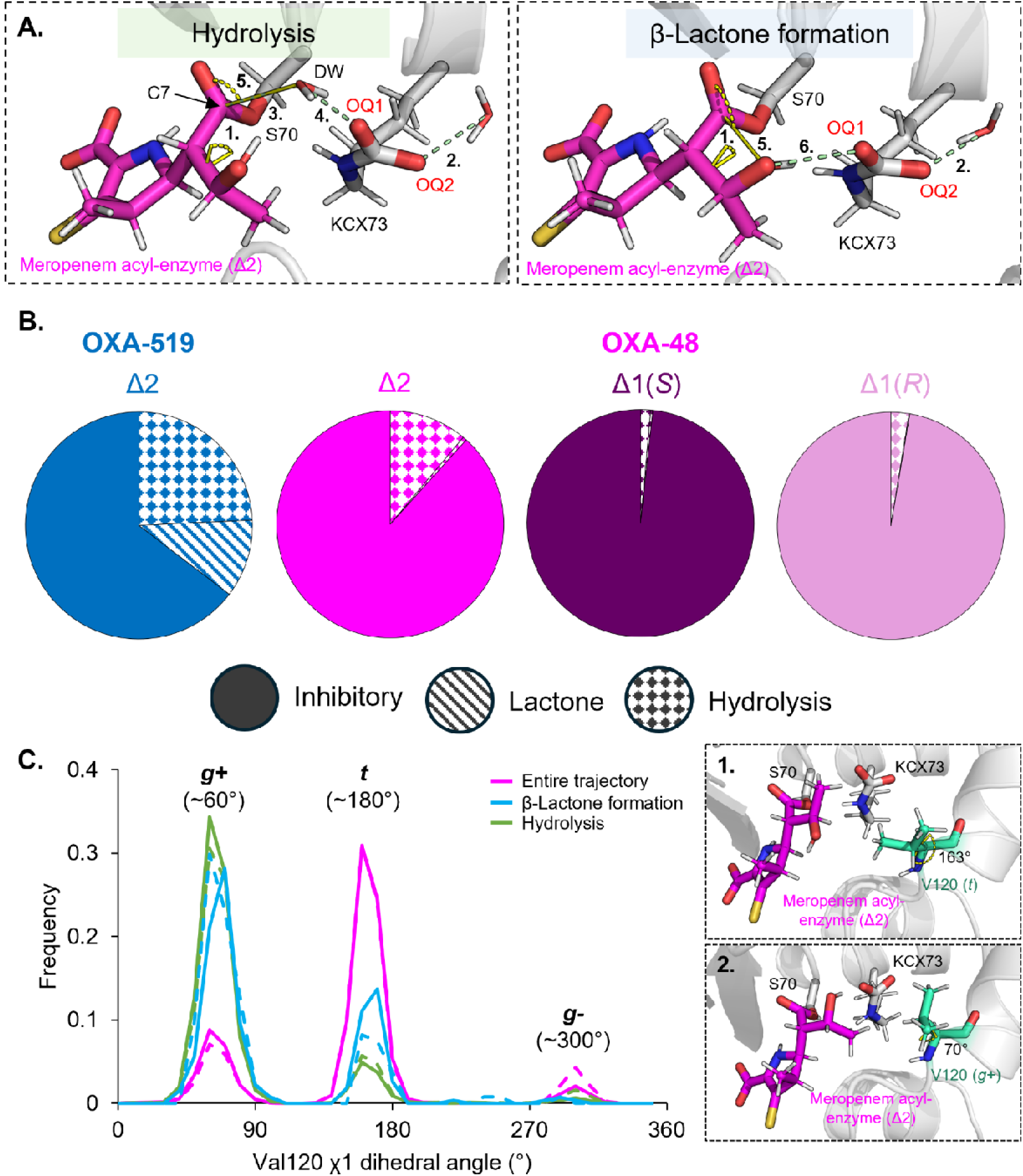
Conformational sampling by 1β-methyl carbapenem-derived acyl-enzymes in MD simulations of OXA-48 and OXA-519. (A) Representative snapshots of MD simulation trajectories of OXA-48:meropenem (Δ2-enamine) in hydrolysis and β-lactone-promoting deacylating conformations. The carbapenem C2 substituent is omitted for clarity. Deacylating conformations were selected by the following parameters: (1.) 6α-hydroxyethyl dihedral angle (C7-C6-C8-O) in conformation I (10 - 90°), (2.) exactly one water (DW) within hydrogen bonding distance (3 Å) of KCX73^OQ^^2^ and a (5.) Bürgi-Dunitz nucleophilic attack angle of (87 - 127°) relative to the C7 carbonyl (6α-hydroxyethyl oxygen for β-lactone formation, DW oxygen for hydrolysis). For hydrolysis, a water (O) is (3.) within 4 Å of the acyl-enzyme C7, (4.) 3 Å of KCX73^OQ^^1^, whilst the β-lactone promoting conformation requires the (6.) acyl-enzyme 6α-hydroxyethyl group to be within hydrogen bonding distance (3 Å) of KCX73^OQ^^1^. (B) Proportions of OXA-48:meropenem (all three tautomers) and OXA-519:meropenem MD simulations where hydrolysis (dots) or β-lactone promoting (lines) conformations are observed, averaged across both active sites. Inhibitory (full colours) refers to neither conformations observed for a given simulation frame. (C) Val120 χ_1_ dihedral angle (N-Cα-Cβ-Cγ1) frequency histogram for OXA-48:meropenem (Δ2) across entire MD trajectories and in frames corresponding to deacylation-promoting acyl-enzyme conformations. Representative MD snapshots where Val120 is in the t (∼180°, panel 1) and g+ (∼60°, panel 2) χ_1_ dihedral rotamer forms are shown in the adjacent inset panels.

Overall, this analysis shows the OXA-519 acyl-enzymes to adopt hydrolysis- and lactone-promoting conformations much more frequently than do those of OXA-48 in MD simulations with both carbapenems, with over a third of the total trajectories in either conformation for OXA-519 when averaged across both active site complexes (**Figs. 6B, S18**). In contrast, for OXA-48 less than a quarter of the simulation trajectories, for any of the three tautomers studied, show the meropenem and ertapenem-derived acyl-enzymes in deacylation-competent binding poses, with the Δ1-imine forms sampling these conformations less frequently than the Δ2-enamine. However, although sampling of acyl-enzyme conformations supporting hydrolysis is favoured in OXA-519, compared to OXA-48, water molecules in a possible deacylating position have markedly reduced survival times (**Fig. S9**). Therefore, although this deacylation-competent conformation is more frequently sampled in OXA-519 acyl-enzymes, it is also likely to be more transient.

The β-lactone-promoting acyl-enzyme conformation is sampled much more frequently in OXA-519 than OXA-48, where it was rarely observed. This is attributed to the 6α-hydroxyethyl oxygen atoms of the ertapenem and meropenem-derived acyl-enzymes generally being positioned closer to KCX73^OQ^^1^ in OXA-519 than in OXA-48 (**Fig. S19**, parameter 1). This is despite conformation I (C7-C6-C8-O dihedral angle of 10 - 90°, likely the most permissive conformation for approach of the 6α-hydroxyethyl oxygen to KCX73^OQ^^1^), being sampled at similar frequencies for the Δ2-enamine tautomers of the meropenem and ertapenem-derived acyl-enzymes of OXA-48 and OXA-519 (**Fig. S20,** parameter 1). We also observed the presence of “mixed” conformations where both hydrolysis and lactone-promoting poses are observed simultaneously, with a water molecule in a deacylating position and the 6α-hydroxyethyl oxygen positioned proximally to KCX73^OQ^^1^ (**Fig. S21**). However, such conformations are generally sampled very rarely, i.e. <1% of all acyl-enzyme simulation trajectories combined. As expected, OXA-519 acyl-enzymes sample this “mixed” conformation more frequently than do those of OXA-48, due to their propensity to more readily adopt the β-lactone-promoting conformation (**Figs. 6B, S18**).

MD trajectories were also analysed to investigate the relationship between the acyl-enzyme geometry and the orientation of the side chain at position 120. Inspection of simulation frames that capture deacylation-competent positions shows that with OXA-48, Val120 more frequently adopts a *g*+ rotamer, despite the *t* rotamer being sampled more often across the entire trajectories (**Fig. 6C**). Crystal structures of uncomplexed OXA-48 also show that the *t* rotamer is preferred (**Fig. S13C**). When Val120 is in the *g*+ rotamer, >95% of simulation frames show the second closest water molecule to KCX73^OQ^^2^ to be out of hydrogen bonding distance (3 Å), whereas <7% of frames satisfy the same criteria for the *t* rotamer (**Fig. S22**). This suggests that rotation of the Val120 side chain in OXA-48 controls hydration around KCX73^OQ^^2^, with the less frequently sampled *g*+ rotamer required to expel a second water molecule from one of the carboxylate oxygens. The presence of this second water molecule close to KCX73 has been previously proposed to increase the energetic barrier for hydrolytic deacylation of meropenem and imipenem by OXA-48, attributed to a decrease in nucleophilicity of the KCX73 general base^23^. Therefore, the Val120 χ_1_ *t* and *g*-rotamers are also likely to be inhibitory to deacylation, whilst rotation to the *g*+ rotamer enables resolution of carbapenem-derived OXA-48 acyl-enzymes. In contrast, during MD simulations of OXA-519, Leu120 remains in the same χ_1_ rotamer in both inhibitory and deacylation-competent poses of bound acyl-enzymes (**Fig. S23**). Indeed, approach of a second water molecule to KCX73^OQ^^2^ is generally impeded in OXA-519, compared to OXA-48 (**Fig. S19**, parameter 4).

## Discussion

Understanding how OXA-48 inactivates carbapenem substrates will help inform the development of new antibiotics or, potentially, inhibitory strategies to overcome OXA-48-mediated resistance.

Crystal structures of OXA-48 complexed with carbapenem-derived acyl-enzymes have been previously-reported and, in general, show similar binding poses to the structures presented here^13,21,22,24–26^. However, structures of 1β-methyl carbapenem-derived acyl-enzymes in the Δ1(*R*)-enantiomer configuration have not been previously reported. Our structure of the OXA-48:ertapenem acyl-enzyme obtained after 1 hour incubation reveals the Δ1(*R*) configuration in dual occupancy with the Δ1(*S*) form, with Lys73 being fully carboxylated (**Fig. 2**). The additional crystal structures presented here also indicate rotation of the carbapenem-derived 6α-hydroxyethyl group, which samples a conformation (conformation III, ∼290° C7-C6-C8-O dihedral angle) not previously seen in crystal structures of OXA-48:carbapenem acyl-enzymes (excluding those in an inactive, “flipped” intermediate^24^), as well as the previously observed conformation II (∼180°) (**Fig 2**). Conformation I (∼50°), described in prior molecular simulations, has yet however to be observed by crystallography in carbapenem-derived acyl-enzymes of OXA-48, consistent with previous QM/MM calculations that suggest this to be the deacylation-competent conformation^23^. This differs from the class A SBL acyl-enzymes, where conformation II of the 6α-hydroxyethyl group was shown by the same simulation methodology to be preferentially deacylated, potentially highlighting a difference between mechanisms of carbapenem hydrolysis in enzymes of classes A and D^33^.

MD simulations of meropenem- and ertapenem-derived acyl-enzymes of OXA-48 in all three possible tautomeric configurations show the Δ2-enamine form to have reduced mobility, and to adopt deacylation-promoting binding poses more frequently than do either of the Δ1-imine forms. This therefore suggests that the Δ2-enamine tautomers of meropenem- and ertapenem-derived OXA-48 acyl-enzymes are likely to be the deacylation-competent forms. Indeed, in all known structures of carbapenem-derived OXA-48 complexes the Δ2 configuration is only observed when Lys73 (which is critical for deacylation) is either mutated, fully decarboxylated or partially decarboxylated, in which case the Δ2-enamine is present in dual occupancy with an alternate tautomer (**Table S6**)^7,9^.

The hydrolytic activity of OXA-48 may therefore derive from its ability to stabilise the Δ2-enamine tautomer of the carbapenem-derived acyl-enzyme, whilst destabilising the alternative Δ1-imine in both its (*R*)- and (*S*)- diastereomeric forms. In fact, our time-course of crystal structures of OXA-519, which compared to OXA-48 has enhanced activity against these carbapenems, captures the Δ2-enamine form of the meropenem-derived acyl-enzyme at shorter time points, with subsequent observation of hydrolysed products bound to the active site. It is only at the later time-points that the Δ2-enamine acyl-enzyme is apparent, corresponding, we propose, to an otherwise catalytically-competent state of the carbapenem-derived acyl-enzyme that has been trapped by enzyme inactivation through decarboxylation of Lys73. In the class A carbapenemase KPC-2, the Δ2-enamine tautomer of the acyl-enzyme is the most hydrolytically labile; we consider it probable that this is also the case for class D SBLs^12^.

We also show differences in the dynamical behaviour of the products of different carbapenem degradation pathways (i.e. hydrolysis and β-lactone formation) bound to OXA-48. Our simulations suggest that, compared to the β-lactone products, hydrolysed meropenem is more readily ejected from the OXA-48 active site. Simulations also indicate that hydrolysed carbapenem products can re-bind the active site, as observed crystallographically in the OXA-519 complexes presented here and in previously determined complex structures of other OXA-48-like enzymes (PDB 6ZRG, 6ZW2^32^). In contrast, the propensity of the β-lactone product to remain in the oxyanion hole of OXA-48 is consistent with its ability to re-acylate, and also to inhibit OXA-48^14^. This may explain why intact β-lactone products have yet to be observed crystallographically as complexes with class D carbapenemases, despite the large number of carbapenem-bound structures deposited. Indeed, crystal structures of PBP1b from *Streptococcus pneumoniae*, an acylation-competent β-lactam target, incubated with β-lactone-containing compounds (albeit not derived from carbapenems) show only covalently-bound acyl-enzymes^34^. Therefore, crystallographic capture of a β-lactone-bound class D carbapenemase may require both isolation of carbapenem-derived β-lactone products for soaking experiments and generation of an acylation-deficient mutant enzyme (e.g. *via* a Ser70Ala substitution). Such studies could be key to understanding the mechanistic basis of β-lactone formation in class D carbapenemases.

Additional factors contributing to carbapenem deacylation by OXA-48-like enzymes were also explored through structural comparisons with OXA-519, which has activity specifically enhanced towards 1β-methyl carbapenems. The single Val120Leu substitution in OXA-519 is unusual as the vast majority of natural OXA-48 variants contain mutations on the loops on the edge of the active site, most notably the β5 - β6 loop which has been demonstrated to be a major determinant of OXA-48 carbapenemase activity^7,32^. In contrast, mutations around the catalytic Lys73 are much rarer in class D carbapenemases^35^. As of March 2026, OXA-519 (V120L), OXA-788 (V120L, β5 – β6 loop deletions and substitutions) and OXA-1382 (V120L, R214G) are the only known natural variants of OXA-48 with a mutation in the deacylating water-channel (Beta-Lactamase DataBase, bldb.eu)^36^. We show here that the ertapenem and meropenem-derived acyl-enzymes more readily adopt both hydrolysis- and β-lactone-promoting conformations in OXA-519 compared to OXA-48, therefore conferring higher turnover rate for 1β-methyl-substituted carbapenems for OXA-519. This can be explained by two main factors. First, in OXA-519 ingress to the active site of a second water molecule, that approaches one of the carboxylate oxygens of KCX73, is greatly reduced, likely enhancing activation of the deacylating water molecule for catalysis^23,37^. In OXA-48, blocking the ingress of this second water molecule requires Val120 to rotate to a less-frequently sampled rotamer, and its more frequent presence in the active site impairs activation of the deacylating water by KCX73. Second, in complexes with OXA-519 the carbapenem 6α-hydroxyethyl oxygen is positioned more frequently towards the KCX73 general base, permitting activation for intramolecular recyclisation and resolution of the acyl-enzyme by β-lactone formation^14^. This relates to the increased ability of OXA-519 to promote β-lactone formation^15^. β-Lactone formation may also be favoured over hydrolysis by the reduced retention time of a water molecule in the deacylating position in OXA-519, compared to OXA-48. The equivalent Val-to-Leu amino acid substitution in another OXA carbapenemase, OXA-23, (Val128Leu) similarly promotes β-lactone formation, and increases *k*_cat_ for meropenem hydrolysis (**Table S1**)^15^. Therefore, control of KCX73 and acyl-enzyme dynamics, through modifications to the deacylating water channel, are likely to be general features of class D carbapenemase activity.

Our findings that OXA-48-class β-lactamases exert tautomer-specific conformational control of the 6α-hydroxyethyl group to favour deacylation, as well as restricting hydration around the catalytic base, indicate potential for modifications to the carbapenem scaffold to modulate susceptibility to these enzymes. One strategy would be to change the composition and/or dynamics of the C6 substituent, achieved in the case of NA-1-157, (a C5α-methyl carbapenem where 6α-hydroxyethyl rotation is restricted that is poorly degraded by some class A and D carbapenemases (including OXA-48)^25,38,39^). Another approach would be to modify the carbapenem scaffold to promote acyl-enzyme tautomerisation to the less reactive Δ1-imine form.

In summary, our results identify and highlight multiple factors: acyl-enzyme tautomerisation, dynamics and active site hydration, alongside the existence of the alternative pathway to β-lactone formation, that collectively enable OXA-48-like SBLs to resolve acyl-enzymes of clinically important 1β-methyl carbapenem antibiotics. As new members of the family continue to be identified it can be anticipated that, as we show here for OXA-519, understanding of the importance of these different contributors to activity will continue to emerge through characterising the effects of point mutations.

## Methods

### Recombinant production of proteins, purification and crystallisation

pOPINF-6HisOXA-48_(23-265)_^40^ was used as a template to generate an OXA-48 V120L expression construct (OXA-519) and express wild-type OXA-48. Forward (5’-CCGCGATGAAATATTCAGTTCTGCCTGTTTATCAAGAATTTG-3’) and reverse (5’-CAAATTCTTGATAAACAGGCAGAACTGAATATTTCATCGCGG-3’) primers were used with the QuikChange II XL site-directed mutagenesis kit (Agilent) to generate pOPINF-6HisOXA-519_(23-265)_, following the manufacturer’s instructions. OXA-48 and OXA-519 were expressed and purified as described previously^18^. Both enzymes were also crystallised under similar conditions as identified before (**Table S5**)^18^. Crystals were then either cryoprotected in 20% (v/v) glycerol or directly immersed in liquid nitrogen for cryo-cooling.

### X-ray data collection and processing

X-ray diffraction data were collected (I03 beamline, Diamond Light Source, UK) and processed as previously described (**Tables S2A-C**)^18^. For refinement, our previously determined structure of unliganded OXA-48 (PDB 9H11^18^), from crystals in the same space-group (*P*6_5_22) and grown under similar conditions, was used as a starting model (waters removed) for phasing by Fourier synthesis (Phenix). With OXA-519 structures, after one round of rigid body refinement (phenix.refine), this yielded *F*_o_-*F*_c_ difference density around Val120, which was then replaced with Leu120 (**Fig. S24**). Ligands were fitted into active site *F*_o_-*F*_c_ difference density using Coot^41^, with ligand geometry restraints generated by Grade2^42^. All structures were refined using phenix.refine, with manual rebuilding and local refitting in Coot, until *R*_work_ and *R*_free_ converged. For OXA-48:ertapenem (1 hour), the Δ1(*R*)-ertapenem tautomer was fitted into the chain A active site using Refmac5 (CCP4)^43,44^ and then merged into the original refined PDB file, as this gave a better fit for the C2 sulphur based on visual inspection of the resultant 2*F*_o_*-F*_c_ map. phenix.refine was then used to generate final model statistics.

### Molecular dynamics simulations

Molecular mechanics molecular dynamics (MD) simulations were run of homodimeric OXA-48 and OXA-519 in their uncomplexed forms and as their meropenem and ertapenem-derived acyl-enzymes. Carbapenem acyl-enzyme derivatives were simulated in the Δ2-enamine tautomer for both enzymes, and the Δ1-imine configuration (in both Δ1(*S*) and Δ1(*R*) enantiomers) for OXA-48 only. In all cases the same carbapenem-derived acyl-enzyme and tautomer was modelled into each enzyme active site for each simulation.

For OXA-48, Δ1(*S*)-ertapenem, Δ1(*S*)-meropenem and Δ2-meropenem were taken from acyl-enzyme structures obtained after crystals were soaked for 1 hour, with other tautomers present in multiple occupancy structures removed from the active sites (**Figs. 2, S1**). The C2-substituent of Δ2-ertapenem (structure obtained from a 16 hour soak, C2 substituent not modelled) was added in Coot to the OXA-48 active site in chain A of the crystallographically observed dimer and was subjected to one round of refinement. As the Δ1(*R*) and Δ2 configurations of ertapenem acyl-enzyme were not identified in chain B of OXA-48 (at 1 hour and 16 hours respectively), they were added by superimposing chains using the SSM algorithm in Coot^45^. The Δ1(*R*) tautomer of meropenem was modelled into chain A of the OXA-48 structure obtained after a 2 hour soak using residual *F*_o_-*F*_c_ density which was not modelled in the final structure (**Fig. S25**). Carboxylated Lys73 (KCX73) was also modelled into those starting structures where Lys73 was decarboxylated (OXA-48:meropenem (2 hours) and OXA-48:ertapenem (16 hours)), using unliganded OXA-48 as a reference. For simulations of meropenem and ertapenem-derived acyl-enzymes with OXA-519, starting structures were taken from 30 minute and 22 hour time-point structures, respectively. The 30 minute OXA-519:meropenem structure was used as a reference to model the carboxylated Lys73 into the 22 hour ertapenem complex, removing the associated chloride ion (**Fig. S17**).

To generate carbapenem-derived hydrolysis and β-lactone product structures, successive QM/MM minimisations (at SCC-DFTB2 level of QM theory^46^) on a representative MD snapshot of the Δ2-meropenem-derived OXA-48 acyl-enzyme were performed using the Sander programme in Amber24^47^. The QM region was defined as the full meropenem-derived covalent adduct, including Ser70 side chain, the deacylating water and the carboxylated Lys73 side chain. Successive QM/MM minimisations were then applied to form the expected product (i.e. moving the 6α-hydroxyethyl oxygen and deacylating water oxygens towards the C7 carbon for β-lactone and hydrolysis product generation, respectively). Minimisations were analogous to the putative deacylation reaction to generate representative product complexes. The products were then subjected to 50 ps of QM/MM MD with restraints on the final positions of the moved atoms (restraining the product bond lengths to prevent re-acylation), followed by a further 50 ps unrestrained QM/MM MD to confirm product stability and accuracy of modelled complex. Structures from the 50 ps unrestrained QM/MM simulations were then used as starting conformations for MD product complex simulations.

The protonation states of ionisable residues at pH 7.4 were determined using PropKa webserver (v2.0). All ionisable residues were predicted to be in their standard protonation states (Asp and Glu deprotonated; Lys and Arg protonated), except Lys208, which was modelled in its deprotonated form. All histidine residues were singly protonated on Nε2. Acyl-enzyme, product complex and carboxylated lysine forcefields were initially generated using general Amber forcefield (GAFF2) parameters, implemented in the Amber24 software package, using the AM1-BCC charge derivation. These partial charges were then replaced with RESP charges calculated by the R.E.D webserver^48^ (as done previously for carboxylated lysine and inhibitor-bound covalent complexes with OXA-48-like enzymes^18,36^). For meropenem-derived hydrolysis and β-lactone products, forcefields were also calculated by R.E.D webserver^48^. The ff14SB MM forcefield was used to describe standard protein residues. Systems were then solvated (in a TIP3P truncated-octahedral water box^49^), minimised and equilibrated as described in Hoff *et al.*^18^. Briefly, waters were minimised (100 cycles of steepest descent, 200 cycles of conjugate gradient) followed by all hydrogen and chloride atoms (1000 cycles of steepest descent, 2000 cycles of conjugate gradient). Systems were then heated in the NVT ensemble to 298 K over 20 ps and equilibrated in the NPT ensemble at 298 K and 1 atm of pressure for 500 ps, restraining protein Cα atoms. Three independent 500 ns simulation trajectories were subsequently run in NPT ensemble at 298 K and 1 atm (as previously^18^). Simulation analyses were performed using CPPTRAJ^50^ and MDAnalysis^51^, including the water survival probability script made available at (https://github.com/Michael-Beer/Water-Survival-Probability). Figures of structures were made using open-source PyMOL (Schrödinger).

## Acknowledgements

All simulations were conducted using the facilities of the Advanced Computing Research Centre at the University of Bristol (http://www.bris.ac.uk/acrc/). The authors thank Diamond Light Source (beamline I03, proposal 31440) and the respective beamline scientists, for their support collecting the X-ray diffraction data presented here. J.S. and C.J.S. acknowledge funding from the U.K. Biotechnology and Biological Sciences Research Council (BBSRC, BB/W001187/1). J.F.H was supported by grant MR/W006308/1 for the GW4 BIOMED2 DTP, awarded to the Universities of Bath, Bristol, Cardiff and Exeter from the Medical Research Council (MRC)/UKRI. M.B. was supported by the BBSRC-funded South-West Biosciences Doctoral Training Partnership (BB/T008741/1). C.L.T. thanks the Medical Research Council for support through the grant UKRI330. This work is part of a project that has received funding from the European Research Council under the European Horizon 2020 research and innovation program (PREDACTED Advanced Grant Agreement no. 101021207) to A.J.M. and J.S.

## Supporting Information

Crystallisation experimental conditions, data processing and validation statistics; supplementary images of crystal structures; analyses of molecular simulations.

## Author Contributions

J.F.H. and J.S. conceived the experiments. J.F.H. performed laboratory experiments and crystallographic data collection/processing, with supervision from M.B., P.H. and C.L.T. J.F.H. undertook molecular simulations with training and supervision from M.B. and A.J.M. J.F.H. drafted the manuscript, with revisions by P.H., M.B., C.L.T., J.S., C.J.S., A.J.M and M.W.K. with input from and final approval by all authors.

## Data availability statement

Structure factors and coordinates for all crystal structures presented here have been deposited with the Protein Data Bank (PDB) with the following accession numbers: OXA-48 in complex with meropenem (1 hour soak) (9TSI), OXA-48 in complex with meropenem (2 hour soak) (9TSJ), OXA-48 in complex with ertapenem (1 hour soak) (9TSK), OXA-48 in complex with ertapenem (16 hour soak) (9TSL), uncomplexed OXA-519 (9TS9), OXA-519 in complex with ertapenem (30 minute soak) (9TSA), OXA-519 in complex with ertapenem (2 hour soak) (9TSB), OXA-519 in complex with ertapenem (4 hour soak) (9TSC), OXA-519 in complex with ertapenem (22 hour soak) (9TSD), OXA-519 in complex with meropenem (30 minute soak) (9TSE), OXA-519 in complex with meropenem (2 hour soak) (9TSF), OXA-519 in complex with meropenem (4 hour soak) (9TSG), OXA-519 in complex with meropenem (22 hour soak) (9TSH). MD data (simulation trajectories and input files) will be made freely available at the University of Bristol Research Data Repository (https://data.bris.ac.uk/). Analysis scripts will be made available upon request.

## Abbreviations

MD: molecular dynamics
OXA: oxacillinase
RMSD: root mean-squared deviation
RMSF: root mean-squared fluctuation
SBL: serine β-lactamase

