## Supplementary Figures and Tables for "Acyl-enzyme dynamics, tautomerisation and hydration regulate turnover of carbapenem antibiotics by the OXA-48 β-lactamase"

**Supporting Information**

Contents

Note S1: Summary of meropenem and ertapenem-derived acyl-enzyme C2 substituent interactions with OXA-48 active sites.	4

Table S1: Carbapenem kinetic parameters for OXA-48, OXA-519 (OXA-48 V120L), OXA-23, OXA-23 V128L.	5

Table S2A: OXA-48 X-ray data collection and structure refinement statistics.	6

Table S2B: OXA-519 X-ray data collection and structure refinement statistics.	7

Table S2C: OXA-519 X-ray data collection and structure refinement statistics.	8

Table S3: RMSD (Cα) of crystal structure compared to uncomplexed OXA-48.	9

Table S4A: Carboxylated Lys73 (KCX73) fitting statistics.	10

Table S4B: Carbapenem-derived complex fitting statistics.	11

Table S5: Summary of crystallisation and ligand soaking experiments summaries.	12

Table S6: Tautomeric state, 6α-hydroxyethyl (C7-C6-C8-O) dihedral angle and Lys73 carboxylation status of reported carbapenem-derived acyl-enzyme complexes with OXA-48.	13

Table S7: Closest water analysis for OXA-48:carbapenem complexes.	14

Figure S1: Chain B active site views of OXA-48 following reaction with meropenem and ertapenem at different ligand soaking time points	15

Figure S2: Hydrogen bonding networks between meropenem and ertapenem derived acyl-enzyme complexes with the active site of OXA-48 in Δ2-enamine, Δ1(*S*)-imine and Δ1(*R*)-imine tautomer configurations	16

Figure S3: OXA-48:ertapenem and meropenem-derived acyl-enzymes in Δ1(*S*) show a water ‘wedged’ between the carbapenem C3 carboxylate and the pyrroline nitrogen	17

Figure S4: RMSD plots across MD simulations of OXA48:carbapenem complexes	18

Figure S5: Average residue RMSF for the OXA-48 and OXA-519 carbapenem-derived acyl-enzyme complexes over MD simulation trajectories	19

Figure S6: Per atom root mean square fluctuation (RMSF) of meropenem and ertapenem-derived complexes with OXA-48 for the Δ2-, Δ1(*S*)- and Δ1(*R*)- tautomers during molecular dynamics simulations	20

Figure S7: Hydrogen bonding analysis of core meropenem and ertapenem-derived acyl-enzyme interactions within the active site of OXA-48 during MD simulations	21

Figure S8: Meropenem- and ertapenem-derived OXA-48 and OXA-519 acyl-enzyme 6α-hydroxyethyl dihedral angle during MD simulations	22

Figure S9: Survival probability of the deacylating water over simulations of meropenem and ertapenem-derived acyl-enzyme complexes with OXA-48 and OXA-519.	23

Figure S10: Bürgi-Dunitz angle for hydrolysis and β-lactone nucleophilic attack trajectories over MD simulations of meropenem- and ertapenem-derived acyl-enzyme complexes with OXA-48.	24

Figure S11: Starting conformations of hydrolysed and β-lactone meropenem products for MD simulations.	25

Figure S12: MD simulations of OXA-48 in complex with nascent meropenem-derived hydrolysed and β-lactone products.	26

Figure S13: Overlays of uncomplexed OXA-48 and OXA-519.	27

Figure S14: Comparison of hydrogen bond networks in the active site of the OXA-519 meropenem-derived acyl-enzyme complexes and hydrolysis products.	28

Figure S15: Unliganded and hydrolysed carbapenem-bound OXA-519 oxyanion hole:water interactions.	29

Figure S16: Overlay of structures of class D SBLs in complex with carbapenem and penem hydrolysed products in the initially observed orientation.	30

Figure S17: Time-course crystallography of meropenem and ertapenem complexes with OXA-519.	31

Figure S18: Conformational sampling of ertapenem-derived acyl-enzymes over MD simulations of OXA-48 and OXA-519.	32

Figure S19: Carbapenem acyl-enzyme complex distance parameters used to filter MD simulations for deacylation promoting conformations.	33

Figure S20: Carbapenem acyl-enzyme complex angle parameters used to filter MD simulations for deacylation promoting conformations.	34

Figure S21: Observation of carbapenem-derived acyl-enzymes in “mixed” hydrolysis and β-lactone promoting conformations.	35

Figure S22: Relationship between Val120 rotamer dynamics and water associated with carboxylated Lys73.	36

Figure S23: Position 120 side-chain rotamer conformational sampling during MD simulations of OXA-48 and OXA-519 meropenem and ertapenem-derived acyl-enzymes.	37

Figure S24: Modelling of Leu120 in OXA-519.	38

Figure S25: Attempted modelling of the meropenem Δ1(*R*)-tautomer into the chain A active site of OXA-48 at 2 hours.	39

Note S1: Summary of meropenem and ertapenem-derived acyl-enzyme C2 substituent interactions with OXA-48 active sites.

The Δ1(*S*)-stereoisomer of the ertapenem derived complex, in which the complete C2 substituent was modelled, is consistently within hydrogen bonding distance of Thr104 (Oγ) via the C2 amide nitrogen (**Fig. S2C**, hydrogen bond network 1). Δ1(*S*)-meropenem derived complex is also positioned to make C2-substituent-mediated interactions with Thr104 (Oγ and N), via direct or indirect (water-mediated, W2) hydrogen bonds (**Fig. S2D**, hydrogen bond network 2). The C2 side chain of the Δ1(*S*)- (1 hour soaked complex, chains A and B) and Δ2-forms of the meropenem derived complex is instead more consistently positioned to interact with the opposing face of the OXA-48 active site (towards the α10 helix), as mediated by a water bridge (W1) between Gln251 (Nε) and the meropenem C2 amide oxygen (**Fig. S2B**, hydrogen bond network 3). The C2 group of the Δ1(*R*)-tautomer of the ertapenem derived complex similarly interacts with Gln251 (Nε and Oε), via the C2 benzoate carboxylate oxygen, which bridges to Gln251 using a water molecule (W3) that is also stabilised by interaction with the side-chain nitrogen of Arg250 (Nη2) and the backbone carbonyl oxygen of Leu247 (**Fig. S2E**, hydrogen bond network 4).

|  | ***K*_M_ (µM)** | | | | ***k*_cat_ (s^-1^)** | | | | ***k*_cat_/*K*_M_ (mM^-1^/s^-1^)** | | | |
| --- | --- | --- | --- | --- | --- | --- | --- | --- | --- | --- | --- | --- |
| β-Lactam | OXA-48 | OXA-519 | OXA-23 | OXA-23 V120L | OXA-48 | OXA-519 | OXA-23 | OXA-23 V120L | OXA-48 | OXA-519 | OXA-23 | OXA-23 V120L |
| Imipenem | 13 | 982 | 5.9 | 5 | 4.8 | 2.1 | 0.59 | 0.01 | 369.2 | 2.1 | 100.0 | 1.9 |
| Meropenem | 11 | 358 | 3.4 | 8.2 | 0.07 | 3.4 | 0.06 | 0.14 | 6.4 | 9.5 | 18.5 | 17.1 |
| Ertapenem | 100 | 83 | 0.5 | - | 0.13 | 1.1 | 0.02 | - | 1.3 | 13.3 | 42.0 | - |

Table S1: Carbapenem kinetic parameters for OXA-48, OXA-519 (OXA-48 V120L), OXA-23, OXA-23 V128L. *Kinetics data for OXA-48 and OXA-519 are from Dabos* et al.^1^*, OXA-23 and OXA-23 V128L with meropenem and imipenem data from Aertker* et al.^2^*, and OXA-23 with ertapenem data from Smith* et al.^3^ *Unknown kinetic parameters are shown as dashes.*

Table S2A: OXA-48 X-ray data collection and structure refinement statistics. *Experiments were performed at beamline I03 of the Diamond Light Source (Didcot, UK). Outer shell statistics are in brackets.*

|  | **OXA-48:**  **ertapenem**  **(1 hour)** | **OXA-48:**  **ertapenem**  **(16 hour)** | **OXA-48:**  **meropenem**  **(1 hour)** | **OXA-48:**  **meropenem**  **(2 hour)** |
| --- | --- | --- | --- | --- |
| **PDB ID** | 9TSK | 9TSL | 9TSI | 9TSJ |
| **Data collection** |  |  |  |  |
| Space group | *P*6_5_22 | *P*6_5_22 | *P*6_5_22 | *P*6_5_22 |
| Molecules/ASU | 2 | 2 | 2 | 2 |
| *Cell dimensions* |  |  |  |  |
| a, b, c (Å) | 122.03, 122.03, 160.14 | 121.98, 121.98, 160.53 | 122.01, 122.01, 160.42 | 122.08, 122.08, 160.46 |
| α, β, γ (°) | 90, 90, 120 | 90, 90, 120 | 90, 90, 120 | 90, 90, 120 |
| Wavelength (Å) | 0.9763 | 0.9763 | 0.9763 | 0.9763 |
| Resolution (Å) | 88.20 - 1.57  (1.60 - 1.57) | 50.17 - 1.51  (1.54 - 1.51) | 88.24 - 1.57  (1.60 - 1.57) | 63.91 - 1.29  (1.31 - 1.29) |
| *R*_pim_ (*I*) | 0.058 (1.292) | 0.044 (1.182) | 0.045 (1.026) | 0.021 (0.905) |
| CC_1/2_ | 0.998 (0.317) | 0.998 (0.547) | 0.999 (0.302) | 1.000 (0.366) |
| *I*/σ (*I*) | 8.7 (0.5) | 9.8 (0.7) | 11.1 (0.7) | 14.6 (0.3) |
| Completeness (%) | 100 (100) | 100 (99.8) | 100 (100) | 100 (99.8) |
| Redundancy | 39.9 (41.2) | 38.2 (31.3) | 39.8 (41.0) | 40.0 (40.1) |
| **Refinement** |  |  |  |  |
| Resolution (Å) | 63.82 - 1.57  (1.63 - 1.57) | 50.17 - 1.52  (1.57 - 1.52) | 63.89 - 1.57  (1.63 - 1.57) | 63.91 - 1.29  (1.34 - 1.29) |
| No. reflections | 97,606 | 107,217 | 98,096 | 174,867 |
| *R*_work_ / *R*_free_ | 0.191 / 0.220 | 0.197 / 0.215 | 0.185 / 0.208 | 0.179 / 0.197 |
| *No. atoms* |  |  |  |  |
| Protein | 4131 | 4131 | 4116 | 4111 |
| Solvent | 502 | 376 | 582 | 475 |
| Ligand | 99 | 32 | 104 | 78 |
| *B-factors (Å^2^)* |  |  |  |  |
| Protein | 25.0 | 24.6 | 22.9 | 26.5 |
| Solvent | 36.9 | 35.0 | 35.9 | 37.7 |
| Ligand | 35.9 | 36.5 | 31.0 | 32.0 |
| *RMS deviations* |  |  |  |  |
| Bond angles (°) | 0.88 | 0.80 | 0.87 | 0.83 |
| Bond lengths (Å) | 0.0062 | 0.0062 | 0.0066 | 0.0061 |
| *Ramachandran (%)* |  |  |  |  |
| Outliers | 0 | 0 | 0 | 0 |
| Favoured | 97.71 | 97.74 | 97.71 | 98.15 |

Table S2B: OXA-519 X-ray data collection and structure refinement statistics. *Experiments were performed at beamline I03 of the Diamond Light Source (Didcot, UK). Outer shell statistics are in brackets.*

|  | **Uncomplexed OXA-519** | **OXA-519:**  **ertapenem**  **(30 mins)** | **OXA-519:**  **ertapenem**  **(2 hour)** | **OXA-519:**  **ertapenem**  **(4 hour)** | **OXA-519:**  **ertapenem**  **(22 hour)** |
| --- | --- | --- | --- | --- | --- |
| **PDB ID** | 9TS9 | 9TSA | 9TSB | 9TSC | 9TSD |
| **Data collection** |  |  |  |  |  |
| Space group | *P*6_5_22 | *P*6_5_22 | *P*6_5_22 | *P*6_5_22 | *P*6_5_22 |
| Molecules/ASU | 2 | 2 | 2 | 2 | 2 |
| *Cell dimensions* |  |  |  |  |  |
| a, b, c (Å) | 122.10, 122.10, 160.19 | 122.04, 122.04, 160.92 | 121.85, 121.85, 160.92 | 122.25, 122.25, 161.00 | 122.08, 122.08, 160.26 |
| α, β, γ (°) | 90, 90, 120 | 90, 90, 120 | 90, 90, 120 | 90, 90, 120 | 90, 90, 120 |
| Wavelength (Å) | 0.9763 | 0.9763 | 0.9763 | 0.9763 | 0.9763 |
| Resolution (Å) | 53.40 - 1.64 (1.67 - 1.64) | 52.84 - 1.83 (1.87 - 1.83) | 53.64 - 1.78 (1.82 - 1.78) | 52.93 - 1.64 (1.67 - 1.64) | 52.86 - 1.80 (1.84 - 1.80) |
| *R*_pim_ (*I*) | 0.060 (1.183) | 0.133 (1.286) | 0.116 (1.831) | 0.081 (1.094) | 0.079 (0.971) |
| CC_1/2_ | 0.998 (0.300) | 0.990 (0.314) | 0.994 (0.316) | 0.997 (0.305) | 0.996 (0.338) |
| *I*/σ (*I*) | 7.1 (0.3) | 5.3 (0.7) | 8.1 (0.6) | 7.2 (0.8) | 8.3 (0.8) |
| Completeness (%) | 100 (99.5) | 100 (100) | 100 (100) | 100 (100) | 99.9 (100) |
| Redundancy | 40.5 (41.7) | 39.4 (39.6) | 39.9 (39.9) | 40.0 (41.2) | 39.4 (39.9) |
| **Refinement** |  |  |  |  |  |
| Resolution (Å) | 50.21 - 1.64 (1.70 - 1.64) | 52.84 - 1.83 (1.90 - 1.83) | 48.57 - 1.78 (1.84 - 1.78) | 52.94 - 1.64 (1.70 - 1.64) | 52.86 - 1.80 (1.86 - 1.80) |
| No. reflections | 85,610 | 62,667 | 67,830 | 86,958 | 65,513 |
| *R*_work_ / *R*_free_ | 0.175 / 0.207 | 0.188 / 0.218 | 0.186 / 0.213 | 0.176 / 0.190 | 0.173 / 0.206 |
| *No. atoms* |  |  |  |  |  |
| Protein | 4128 | 4123 | 4126 | 4160 | 4077 |
| Solvent | 625 | 421 | 438 | 492 | 441 |
| Ligand | - | 102 | 68 | 102 | 66 |
| *B-factors (Å^2^)* |  |  |  |  |  |
| Protein | 28.0 | 19.7 | 22.2 | 21.2 | 23.4 |
| Solvent | 39.5 | 31.2 | 34.6 | 35.8 | 35.8 |
| Ligand | - | 26.0 | 36.1 | 29.3 | 38.2 |
| *RMS deviations* |  |  |  |  |  |
| Bond angles (°) | 0.77 | 0.84 | 0.85 | 0.85 | 0.83 |
| Bond lengths (Å) | 0.0062 | 0.0075 | 0.0070 | 0.0079 | 0.0070 |
| *Ramachandran (%)* |  |  |  |  |  |
| Outliers | 0 | 0 | 0 | 0 | 0 |
| Favoured | 97.71 | 98.54 | 97.71 | 97.92 | 98.56 |

Table S2C: OXA-519 X-ray data collection and structure refinement statistics. *Experiments were performed at beamline I03 of the Diamond Light Source (Didcot, UK). Outer shell statistics are in brackets.*

|  | **OXA-519:**  **meropenem**  **(30 mins)** | **OXA-519:**  **meropenem**  **(2 hour)** | **OXA-519:**  **meropenem**  **(4 hour)** | **OXA-519:**  **meropenem**  **(22 hour)** |
| --- | --- | --- | --- | --- |
| **PDB ID** | 9TSE | 9TSF | 9TSG | 9TSH |
| **Data collection** |  |  |  |  |
| Space group | *P*6_5_22 | *P*6_5_22 | *P*6_5_22 | *P*6_5_22 |
| Molecules/ASU | 2 | 2 | 2 | 2 |
| *Cell dimensions* |  |  |  |  |
| a, b, c (Å) | 122.21, 122.21, 160.76 | 122.23, 122.23, 160.51 | 122.17, 122.17, 161.22 | 122.02, 122.02, 160.39 |
| α, β, γ (°) | 90, 90, 120 | 90, 90, 120 | 90, 90, 120 | 90, 90, 120 |
| Wavelength (Å) | 0.9763 | 0.9763 | 0.9763 | 0.9763 |
| Resolution (Å) | 53.59 - 1.95  (2.00 - 1.95) | 52.93 - 1.72  (1.75 - 1.72) | 50.26 - 1.92  (1.95 - 1.92) | 53.46 - 1.43  (1.46 - 1.43) |
| *R*_pim_ (*I*) | 0.158 (1.793) | 0.067 (1.617) | 0.276 (0.693) | 0.032 (0.931) |
| CC_1/2_ | 0.984 (0.320) | 0.998 (0.339) | 0.947 (0.342) | 0.999 (0.313) |
| *I*/σ (*I*) | 4.7 (0.5) | 9.9 (0.6) | 4.2 (0.5) | 11.9 (0.4) |
| Completeness (%) | 100 (100) | 100 (100) | 100 (100) | 100 (99.7) |
| Redundancy | 38.9 (37.5) | 39.2 (40.3) | 35.5 (34.9) | 40.0 (39.3) |
| **Refinement** |  |  |  |  |
| Resolution (Å) | 48.65 - 1.95  (2.02 - 1.95) | 52.93 - 1.72  (1.78 - 1.72) | 50.26 - 1.92  (1.99 - 1.92) | 52.84 - 1.43  (1.48 - 1.43) |
| No. reflections | 51,635 | 75,229 | 54,905 | 128,965 |
| *R*_work_ / *R*_free_ | 0.204 / 0.247 | 0.180 / 0.204 | 0.188 / 0.224 | 0.176 / 0.186 |
| *No. atoms* |  |  |  |  |
| Protein | 4127 | 4113 | 4113 | 4215 |
| Solvent | 318 | 399 | 373 | 456 |
| Ligand | 52 | 34 | 52 | 52 |
| *B-factors (Å^2^)* |  |  |  |  |
| Protein | 24.9 | 25.0 | 31.3 | 23.0 |
| Solvent | 34.2 | 37.2 | 41.7 | 36.7 |
| Ligand | 40.9 | 33.8 | 42.8 | 36.0 |
| *RMS deviations* |  |  |  |  |
| Bond angles (°) | 0.85 | 0.86 | 0.82 | 0.83 |
| Bond lengths (Å) | 0.0076 | 0.0073 | 0.0069 | 0.0064 |
| *Ramachandran (%)* |  |  |  |  |
| Outliers | 0 | 0 | 0 | 0 |
| Favoured | 98.12 | 97.71 | 97.92 | 98.15 |

Table S3: RMSD (Cα) of crystal structure compared to uncomplexed OXA-48. *Crystal structures (including both chains in the asymmetric unit) were aligned to uncomplexed OXA-48 (PDB 9H11), previously determined under similar conditions*^4^*. Alignments and RMSD were calculated using PyMOL (Schrödinger).*

| **Crystal structure** | **PDB ID** | **# of residues aligned** | **Cα RMSD (Å)** |
| --- | --- | --- | --- |
| OXA-48:ertapenem (1 hour) | 9TSK | 479 | 0.19 |
| OXA48:ertapenem (16 hour) | 9TSL | 481 | 0.31 |
| OXA-48:meropenem (1 hour) | 9TSI | 475 | 0.20 |
| OXA-48:meropenem (2 hour) | 9TSJ | 484 | 0.28 |
| Uncomplexed OXA-519 | 9TS9 | 486 | 0.18 |
| OXA-519:ertapenem (30 mins) | 9TSA | 469 | 0.26 |
| OXA-519:ertapenem (2 hour) | 9TSB | 464 | 0.26 |
| OXA-519:ertapenem (4 hour) | 9TSC | 464 | 0.25 |
| OXA-519:ertapenem (22 hour) | 9TSD | 488 | 0.28 |
| OXA-519:meropenem (30 mins) | 9TSE | 477 | 0.26 |
| OXA-519:meropenem (2 hour) | 9TSF | 470 | 0.28 |
| OXA-519:meropenem (4 hour) | 9TSG | 469 | 0.27 |
| OXA-519:meropenem (22 hour) | 9TSH | 489 | 0.30 |

Table S4A: Carboxylated Lys73 (KCX73) fitting statistics. *(A) Carboxylated Lys73 (KCX73) occupancy and RSCC fitting statistics, partial occupancy KCX73 was refined in dual-occupancy with a decarboxylated, free lysine to a total occupancy of 1. RSCC values were calculated by the PDB validation server.*

| **Crystal structure** | **PDB ID** | **KCX73 occupancy** | | **KCX73 RSCC** | |
| --- | --- | --- | --- | --- | --- |
|  |  | Chain A | Chain B | Chain A | Chain B |
| OXA-48:ertapenem (1 hour) | 9TSK | 1 | 0.63 | 0.97 | 0.97 |
| OXA48:ertapenem (16 hour) | 9TSL | - | - | - | - |
| OXA-48:meropenem (1 hour) | 9TSI | 0.49 | 0.39 | 0.97 | 0.97 |
| OXA-48:meropenem (2 hour) | 9TSJ | - | - | - | - |
| Uncomplexed OXA-519 | 9TS9 | 1 | 1 | 0.98 | 0.97 |
| OXA-519:ertapenem (30 mins) | 9TSA | 1 | 1 | 0.97 | 0.97 |
| OXA-519:ertapenem (2 hour) | 9TSB | 1 | 1 | 0.96 | 0.97 |
| OXA-519:ertapenem (4 hour) | 9TSC | 1 | 1 | 0.98 | 0.98 |
| OXA-519:ertapenem (22 hour) | 9TSD | - | - | - | - |
| OXA-519:meropenem (30 mins) | 9TSE | 1 | 1 | 0.95 | 0.96 |
| OXA-519:meropenem (2 hour) | 9TSF | 1 | 1 | 0.97 | 0.96 |
| OXA-519:meropenem (4 hour) | 9TSG | 0.44 | 0.51 | 0.95 | 0.94 |
| OXA-519:meropenem (22 hour) | 9TSH | - | - | - | - |

Table S4B: Carbapenem-derived complex fitting statistics. *Occupancy and RSCC values of carbapenem-derived acyl-enzyme and product complexes of OXA-48 and OXA-519. The pyrroline tautomeric state(s) of carbapenem-derived acyl-enzymes for dual-occupancy ligands are stated in parentheses. RSCC values were calculated by the PDB validation server.*

| **Crystal structure** | **PDB ID** | **Ligand occupancy** | | **Ligand RSCC** | |
| --- | --- | --- | --- | --- | --- |
|  |  | Chain A | Chain B | Chain A | Chain B |
| OXA-48:ertapenem (1 hour) | 9TSK | 0.47 (Δ1(*R*)) 0.53 (Δ1(*S*)) | 1 | 0.85 (Δ1(*R*)) 0.80 (Δ1(*S*)) | 0.85 |
| OXA48:ertapenem (16 hour) | 9TSL | 1 | 1 | 0.83 | 0.89 |
| OXA-48:meropenem (1 hour) | 9TSI | 0.71 (Δ2) 0.29 (Δ1(*S*)) | 0.59 (Δ2) 0.41 (Δ1(*S*)) | 0.87 (Δ2) 0.87 (Δ1(*S*)) | 0.91 (Δ2) 0.77 (Δ1(*S*)) |
| OXA-48:meropenem (2 hour) | 9TSJ | 0.53 (Δ2) 0.47 (Δ1(*S*)) | 1 | 0.78 (Δ2) 0.80 (Δ1(*S*)) | 0.90 |
| Uncomplexed OXA-519 | 9TS9 | - | - | - | - |
| OXA-519:ertapenem (30 mins) | 9TSA | 0.55 0.45 | 1 | 0.82 0.82 | 0.91 |
| OXA-519:ertapenem (2 hour) | 9TSB | 0.86 | 0.86 | 0.86 | 0.86 |
| OXA-519:ertapenem (4 hour) | 9TSC | 0.38 0.50 | 0.81 | 0.78 0.86 | 0.92 |
| OXA-519:ertapenem (22 hour) | 9TSD | 1 | 1 | 0.93 | 0.88 |
| OXA-519:meropenem (30 mins) | 9TSE | 1 | 1 | 0.87 | 0.89 |
| OXA-519:meropenem (2 hour) | 9TSF | 0.75 | 0.62 | 0.86 | 0.80 |
| OXA-519:meropenem (4 hour) | 9TSG | 1 | 1 | 0.92 | 0.92 |
| OXA-519:meropenem (22 hour) | 9TSH | 1 | 1 | 0.93 | 0.90 |

Table S5: Summary of crystallisation and ligand soaking experiments summaries.

| Protein | Ligand | Ligand soak time | Ligand concentration | Crystallisation condition |
| --- | --- | --- | --- | --- |
| OXA-48 | Ertapenem | 1 hr | 20 mM | 0.1 M Tris pH 8.8, 50% PEG 400 |
|  | Ertapenem | 16 hr | 20 mM | 0.1 M phosphate pH 7.3, 57.5% PEG 400 |
|  | Meropenem | 1 hr | 10 mM | 0.1 M Tris pH 8.8, 50% PEG 400 |
|  | Meropenem | 2 hr | 10 mM | 0.1 M phosphate pH 7.5, 52.5% PEG 400 |
| OXA-519 | - | - | - | 0.1 M Tris pH 9.0, 50% PEG 400 |
|  | Ertapenem | 30 mins | 10 mM | 0.1 M Tris pH 8.8, 50% PEG 400 |
|  | Ertapenem | 2 hr | 10 mM | 0.1 M Tris pH 8.6, 50% PEG 400 |
|  | Ertapenem | 4 hr | 10 mM | 0.1 M Tris pH 8.8, 50% PEG 400 |
|  | Ertapenem | 22 hr | 1 mM | 0.1 M Tris pH 8.6, 50% PEG 400 |
|  | Meropenem | 30 mins | 10 mM | 0.1 M Tris pH 8.8, 50% PEG 400 |
|  | Meropenem | 2 hr | 10 mM | 0.1 M Tris pH 8.6, 50% PEG 400 |
|  | Meropenem | 4 hr | 10 mM | 0.1 M Tris pH 8.8, 50% PEG 400 |
|  | Meropenem | 22 hr | 10 mM | 0.1 M Tris pH 8.6, 50% PEG 400 |

Table S6: Tautomeric state, 6α-hydroxyethyl (C7-C6-C8-O) dihedral angle and Lys73 carboxylation status of reported carbapenem-derived acyl-enzyme complexes with OXA-48. **OXA-48 K73A mutant, **OXA-48* N*-acetyl-K73 modification, ***An atypical (i.e. “flipped”) binding mode observed for the acyl-enzyme complex (****Fig. 5C-D****)*^5^*.*

| **Carbapenem** | **Tautomer** | **PDB ID** | **Chain** | **C7-C6-C8-O dihedral angle (°)** | **Lys73 carboxylated?** |
| --- | --- | --- | --- | --- | --- |
| Ertapenem | Δ2 | 6P99 | A | 162.3 | No |
|  |  | 9TSL *This work* (16 hr) | A | 177.4 | No |
| Meropenem |  | 6P98 | A | 175.9 | No |
|  |  |  | B | 177.4 | No |
|  |  | 6PT1 | A | 184.1 | No |
|  |  |  | B | 186.0 | No |
|  |  |  | C | 173.3 | No |
|  |  |  | D | 181.7 | No |
|  |  | 7KHQ* | A | 174.5 | - |
|  |  |  | B | 188.3 | - |
|  |  | 9TSI *This work* (1 hr) | A | 191.3 | Partially |
|  |  |  | B | 178.3 | Partially |
|  |  | 9TSJ *This work* (2 hr) | A | 177.3 | No |
| Imipenem |  | 6P97 | A | 176.9 | No |
|  |  |  | B | 174.2 | No |
|  |  | 7KH9* | A | 147.1 | - |
|  |  |  | B | 157.5 | - |
| Doripenem |  | 6P9C | A | 157.6 | No |
|  |  |  | B | 171.6 | No |
| NA-1-157 |  | 8FAJ | A | 170.4 | No |
|  |  |  | B | 169.4 | No |
| Ertapenem | Δ1(*S*) | 6ZRJ | A | 168.9 | Yes |
|  |  |  | B | 183.4 | No |
|  |  |  | C | 173.7 | Yes |
|  |  |  | D | 183.6 | Yes |
|  |  |  | E | 179.5 | Yes |
|  |  |  | F | 168.3 | Yes |
|  |  |  | G | 176.6 | Yes |
|  |  |  | H | 169.7 | Yes |
|  |  | 9TSK *This work* (1 hr) | A | 175.2 | Yes |
|  |  |  | B | 171.0 | Partially |
|  |  | 9TSL *This work* (16 hr) | B | 176.8 | No |
| Meropenem |  | 6ZRP | A | 171.9 | Yes |
|  |  |  | B | 174.3 | No |
|  |  |  | C | 179.7 | No |
|  |  |  | D | 180.7 | Yes |
|  |  | 9TSI *This work* (1 hr) | A | 320.9 | Partially |
|  |  |  | B | 178.6 | Partially |
|  |  | 9TSJ *This work* (2 hr) | A | 182.0 | No |
|  |  |  | B | 176.4 | No |
| Ertapenem | Δ1(*R*) | 9TSK *This work* (1 hr) | A | 279.4 | Yes |
| Imipenem |  | 6PTU | A | 164.5 | No |
|  |  |  | B | 169.5 | No |
|  |  |  | C | 176.4 | No |
|  |  |  | D | 170.2 | No |
|  |  | 7PSF | A | 176.7 | Yes |
|  |  |  | B | 144.8 | Yes |
|  |  | 7PFN** | A | 172.5 | - |
|  |  |  | B*** | 294.8 | - |
|  |  | 7Q14** | A*** | 287.1 | - |
|  |  |  | B*** | 280.3 | - |
|  |  |  | C | 151.2 | - |
|  |  |  | D | 187.7 | - |
|  |  |  | E | 162.5 | - |
|  |  |  | F | 185.0 | - |
|  |  |  | F*** | 129.9 | - |
|  |  |  | G | 181.8 | - |
|  |  |  | G*** | 272.5 | - |
|  |  |  | H*** | 168.2 | - |
| Doripenem |  | 6PXX* | A | 178.7 | - |
|  |  |  | B | 180.3 | - |

Table S7: Closest water analysis for OXA-48:carbapenem complexes. *(A) Proportion of MD simulations of OXA-48:carbapenem derived complexes where the closest water (any atom) to the acyl-enzyme carbonyl carbon is <3 Å away, for each carbapenem 6α-hydroxyethyl conformation (0-III). The 6α-hydroxyethyl conformation was assigned by measuring the C7-C6-C8-O dihedral angle ranges: I = 10 - 90°, II = 140 – 220°, III = 250 – 330°, 0 = any angle not within these ranges.*

| 6α-hydroxyethyl conformation | **Percentage of simulation where the closest water is less than 3 Å** | | | | | |
| --- | --- | --- | --- | --- | --- | --- |
|  | OXA-48:meropenem | | | OXA-48:ertapenem | | |
|  | Δ2 | Δ1(*S*) | Δ1(*R*) | Δ2 | Δ1(*S*) | Δ1(*R*) |
| 0 | 0.0% | 0.1% | 0.1% | 0.0% | 0.1% | 0.1% |
| I | 28.2% | 5.2% | 6.8% | 26.7% | 5.8% | 15.4% |
| II | 0.0% | 2.7% | 0.9% | 0.0% | 2.0% | 0.3% |
| III | 0.0% | 0.5% | 0.3% | 0.0% | 0.2% | 0.4% |

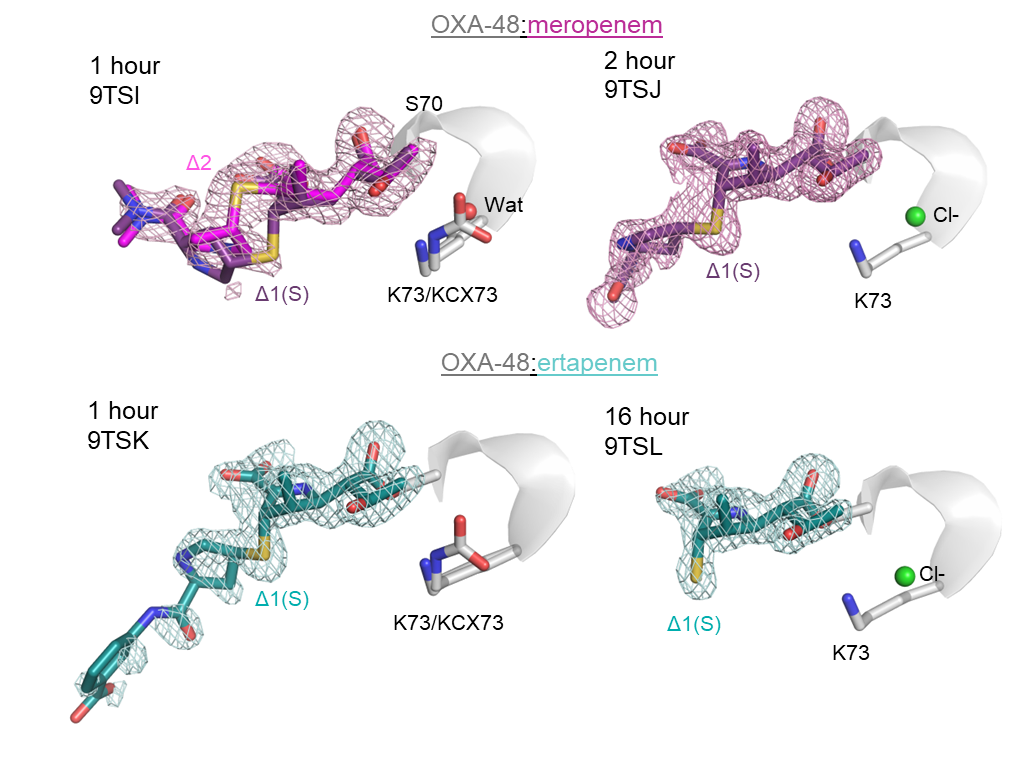
Figure S1: Chain B active site views of OXA-48 following reaction with meropenem and ertapenem at different ligand soaking time points**.** *Lys73 and Ser70 are shown as grey sticks with meropenem-derived acyl-enzymes as pink/purple sticks, and ertapenem-derived complexes as blue sticks.* F*_o_-*F*_c_ ligand omit maps are shown as mesh (contoured to 3σ) and chloride ions and waters proximal to Lys73 as green and red spheres, respectively.*

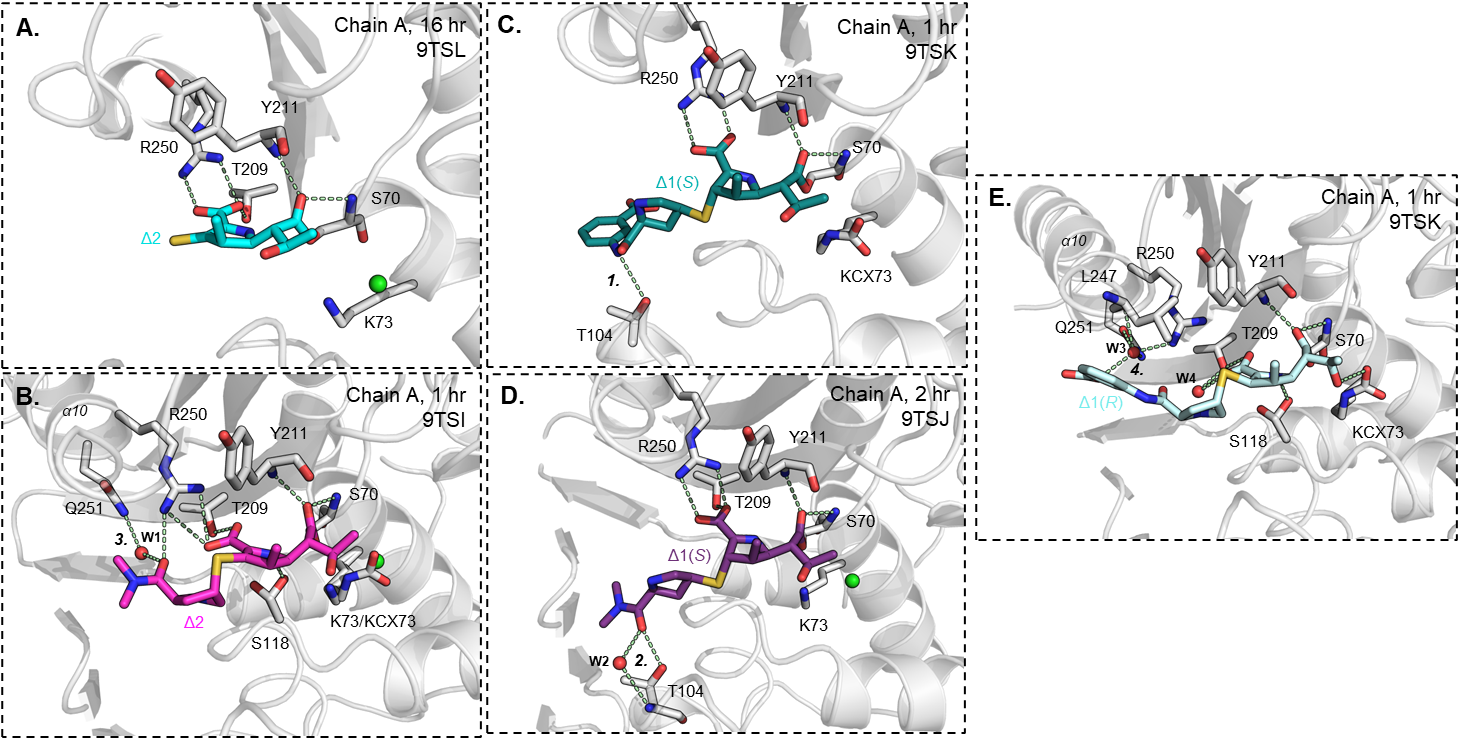
Figure S2: Hydrogen bonding networks between meropenem and ertapenem derived acyl-enzyme complexes with the active site of OXA-48 in Δ2-enamine, Δ1(*S*)-imine and Δ1(*R*)-imine tautomer configurations**.** *The ertapenem- (shades of blue) and meropenem-derived (shades of pink) complexes and interacting OXA-48 active site residues (grey) are represented as sticks.* *Waters (numbered in order of their appearance in the text) and chloride ions are shown as red and green spheres, respectively. Potential hydrogen bonds are shown as pale green dashed lines, with	carbapenem-derived C2 substituent-mediated hydrogen bonding networks numbered 1 to 4 (see* ***Note S1*** *for more details). Ligand soak time and OXA-48 chain is specified in the top right corner of each panel and the modelled carbapenem-derived acyl-enzyme tautomer form is labelled adjacent to the ligand.*

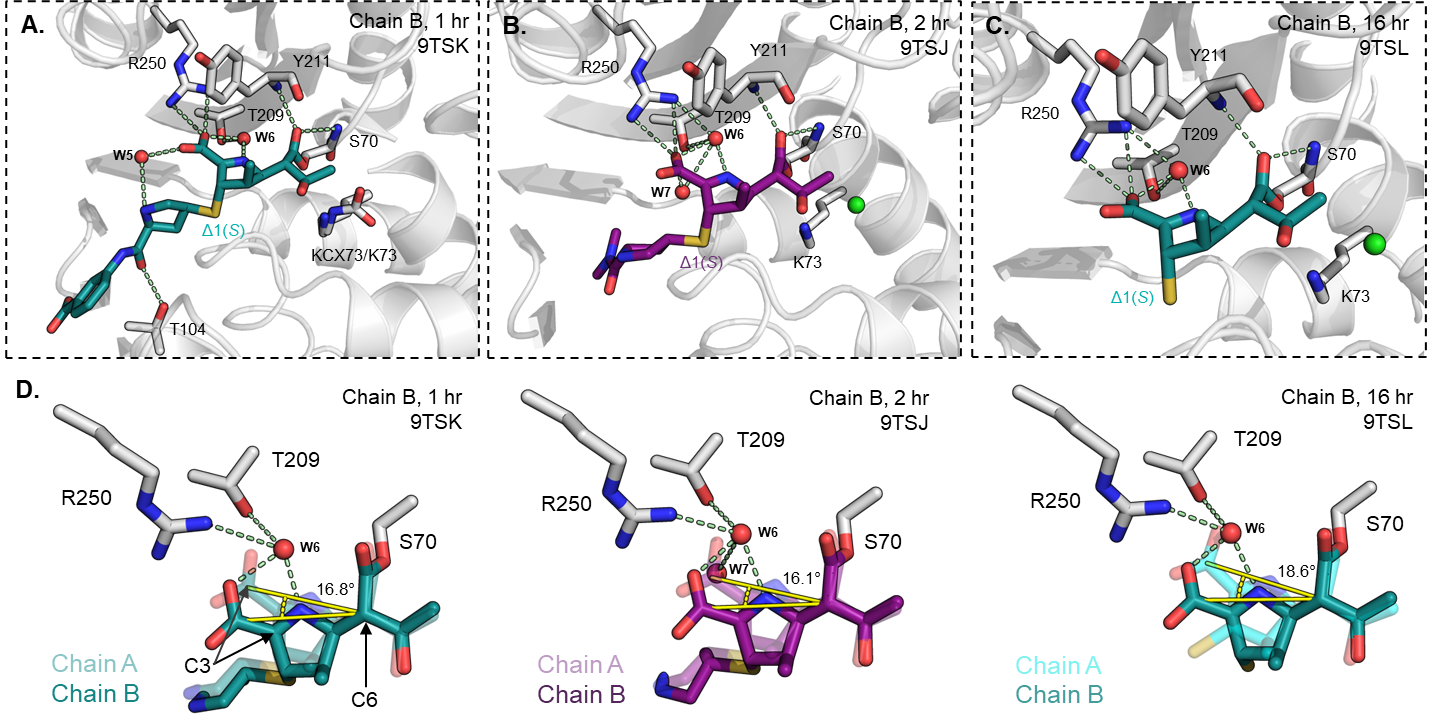
Figure S3: OXA-48:ertapenem and meropenem-derived acyl-enzymes in Δ1(*S*) show a water ‘wedged’ between the carbapenem C3 carboxylate and the pyrroline nitrogen**.** *OXA-48 active site depicted as shown in* ***Fig. S2****. with Δ1(*S*)-configurations of ertapenem and meropenem acyl-enzyme complexes in chain B of OXA-48 at: (A) 1 hour, (B) 2 hour and (C) 16 hour soak times, respectively. (D) Close up of an overlay of the carbapenem acyl-enzyme (transparent) shown above with the opposing active site of OXA-48, where the ‘wedged’ water (W6) is not observed. Possible hydrogen bonding interactions with the ‘wedged’ water are indicated by pale green dashed lines and the angle of rotation between the C3 carboxylate across active sites, relative to the C6 carbon, is in yellow.*

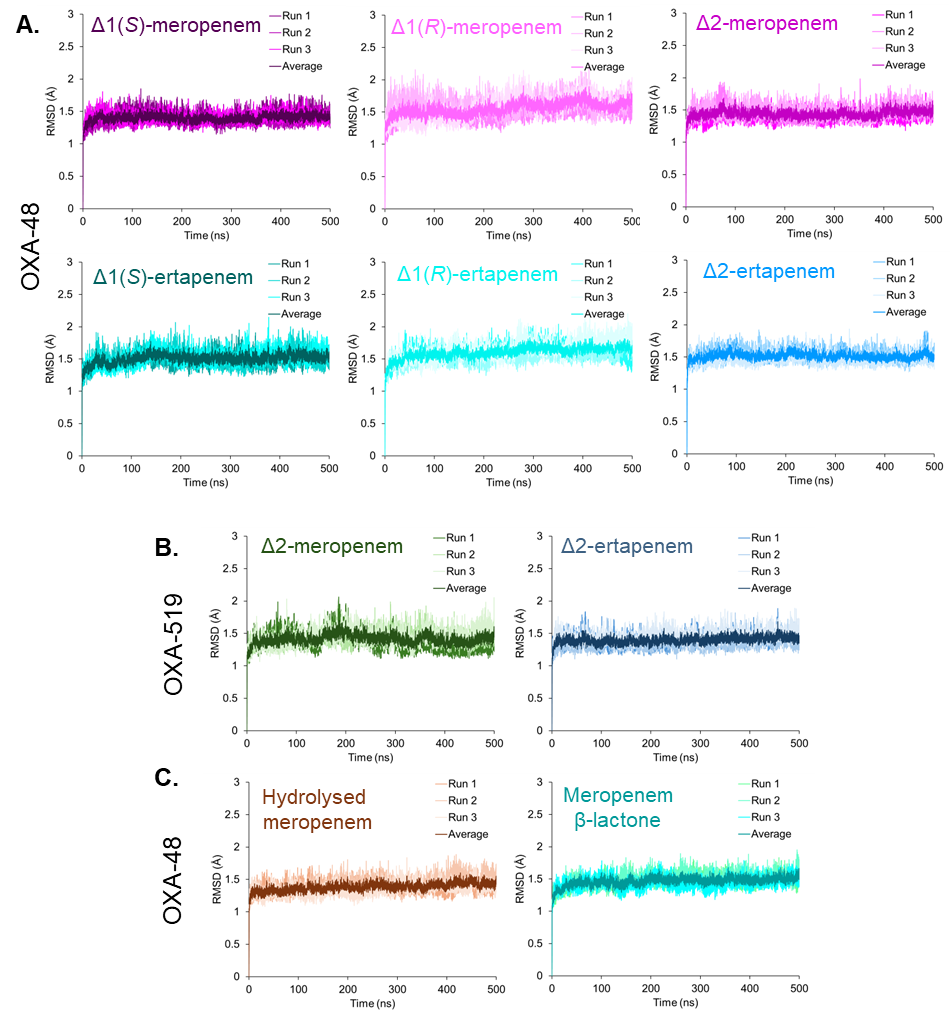

Figure S4: RMSD plots across MD simulations of OXA48:carbapenem complexes**.** *Simulations of meropenem and ertapenem-derived acyl-enzyme complexes with (A) OXA-48 in Δ1-imine and Δ2-enamine tautomer configurations (B) OXA-519 in the Δ2-enamine form. (C) OXA-48:meropenem hydrolysed and β-lactone product complexes. RMSD plots were calculated using all non-hydrogen protein atoms across both chains of OXA-48 relative to the first frame of each simulation trajectory, excluding residues 21-24 that comprise the purification tag/signal sequence.*

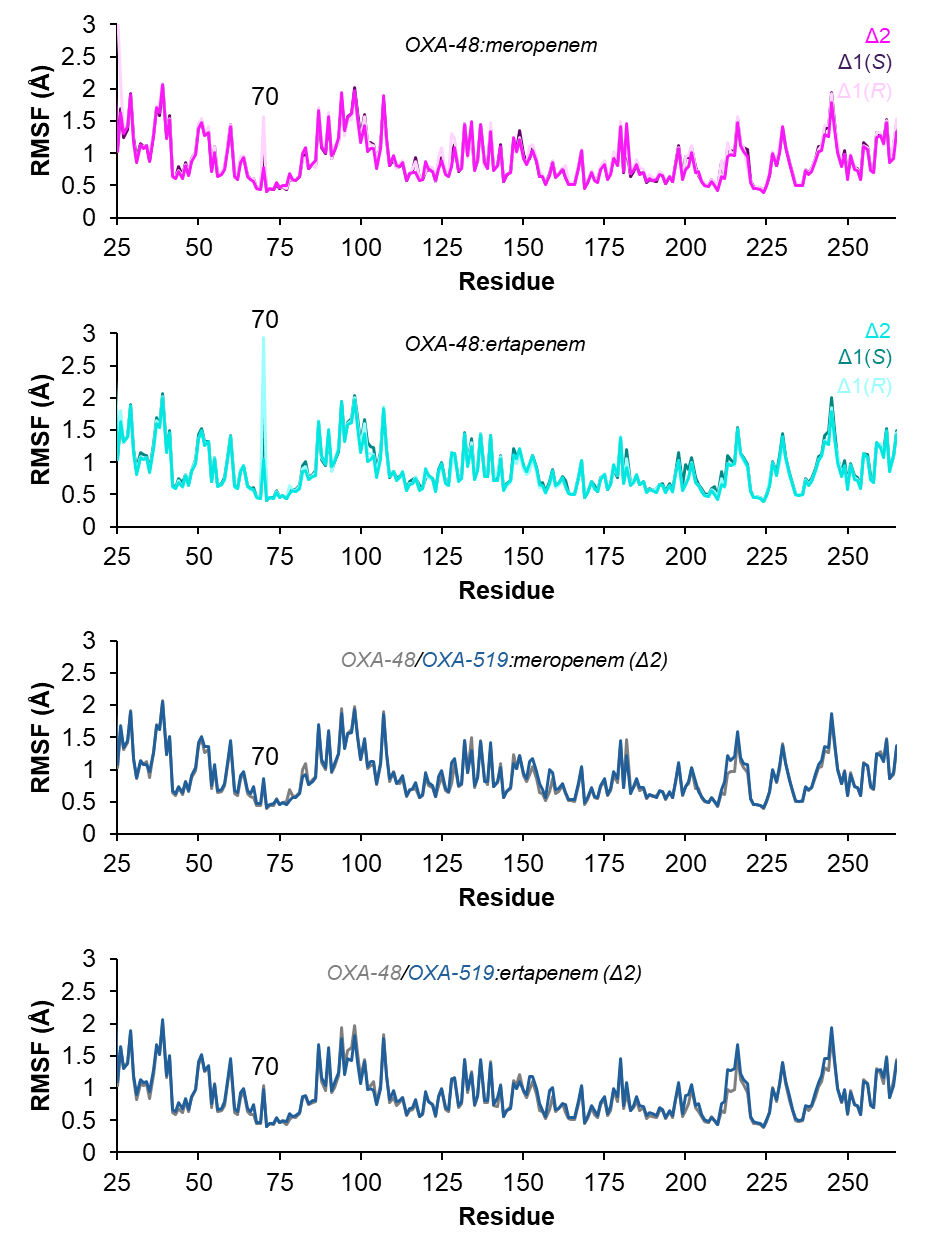
Figure S5: Average residue RMSF for the OXA-48 and OXA-519 carbapenem-derived acyl-enzyme complexes over MD simulation trajectories**.** *RMSF (Å) is calculated using the minimised structure as a reference and averaged over each chain of OXA-48/OXA-519 homodimers. Modified residue 70 composing Ser70 and the carbapenem-derived acyl-enzyme is highlighted.*

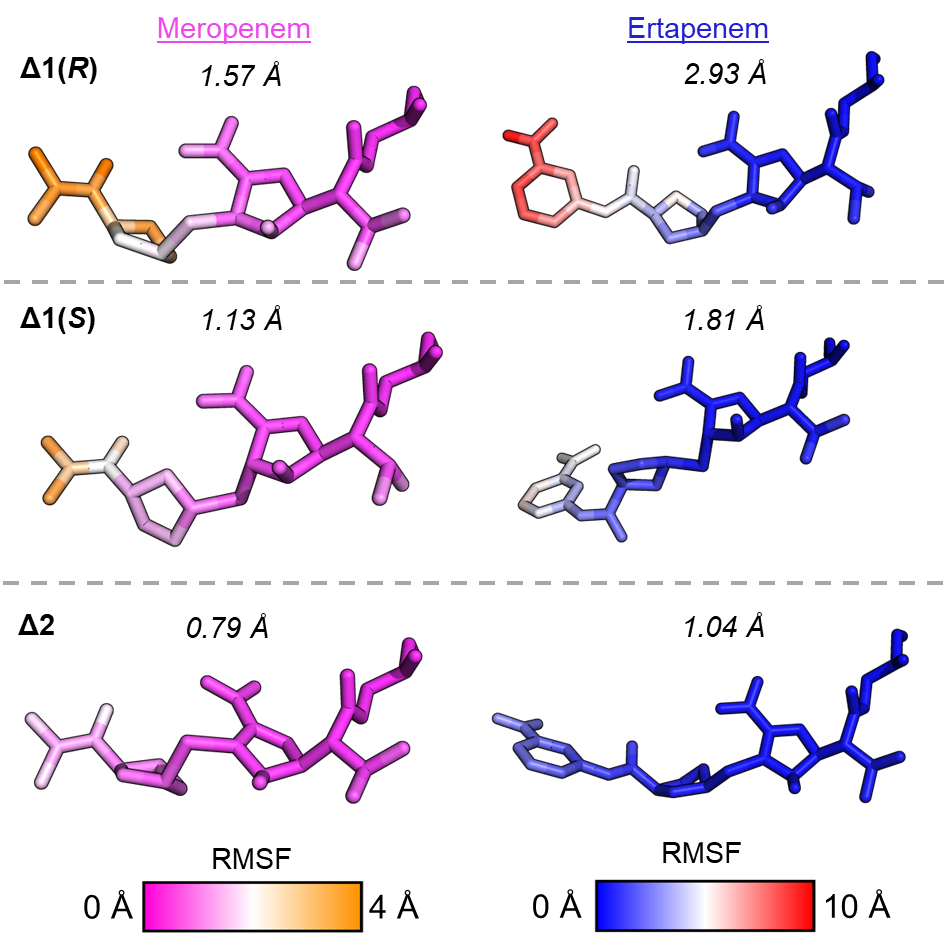

Figure S6: Per atom root mean square fluctuation (RMSF) of meropenem and ertapenem-derived complexes with OXA-48 for the Δ2-, Δ1(*S*)- and Δ1(*R*)- tautomers during molecular dynamics simulations**.** *RMSF per atom (Å), averaged across both active sites over 1.5 µs of simulation time of OXA-48 in complex with the respective carbapenem-derived acyl-enzyme complex, coloured using the scales shown. The average RMSF of the carbapenem:Ser70 complex is shown is in italics above each acyl-enzyme complex.*

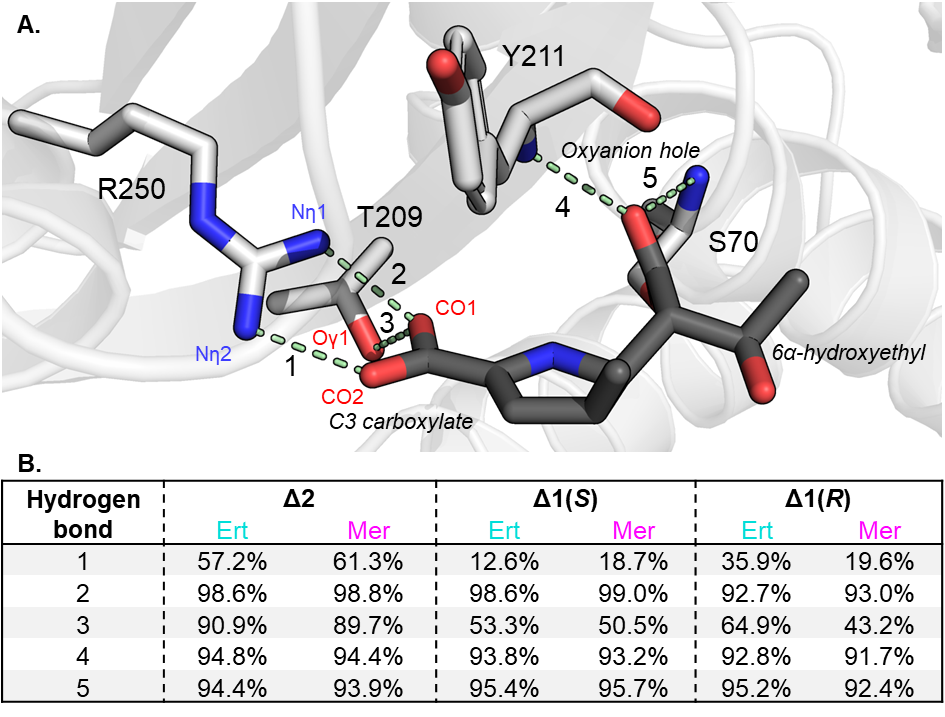
Figure S7: Hydrogen bonding analysis of core meropenem and ertapenem-derived acyl-enzyme interactions within the active site of OXA-48 during MD simulations**.** *(A) Chain A active site (light grey) of the minimised structure of OXA-48 in complex with meropenem in the Δ2 tautomer configuration (dark grey). The meropenem C2-substituent has been removed and components of the core carbapenem scaffold are labelled in italics. Active site residues commonly observed to facilitate carbapenem binding in OXA-48 are shown as light grey sticks, with key hydrogen bonding interactions represented by pale green dashes and numbered 1 to 5. (B) Table showing the proportion of simulations that the hydrogen bonds identified above are observed (3.0 Å heavy atom distance cutoff, 135° hydrogen bond angle cutoff) averaged across both chains of OXA-48 with ertapenem (‘Ert’) and meropenem (‘Mer’) in the various tautomer configurations stated.*

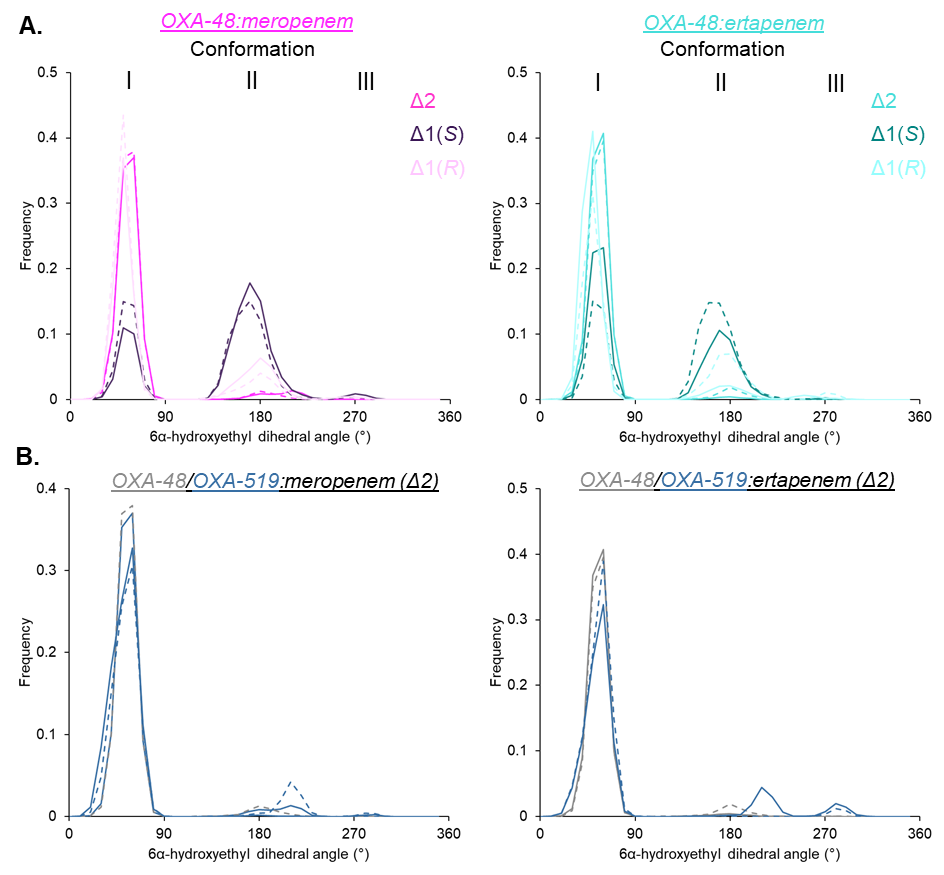

Figure S8: Meropenem- and ertapenem-derived OXA-48 and OXA-519 acyl-enzyme 6α-hydroxyethyl dihedral angle during MD simulations**.** *Frequency histograms of the carbapenem 6α-hydroxyethyl* *C7-C6-C8-O dihedral angle over a 1.5 μs simulation time. (A)* *Meropenem (shades of pink) and ertapenem-derived (shades of turquoise) acyl-enzyme complexes of OXA-48 in the Δ1(*S*)-, Δ1(*R*)-, and Δ2-tautomeric forms. (B) OXA-48 and OXA-519 acyl-enzyme complexes of meropenem and ertapenem in Δ2-enamine forms. Dihedral angles were separated into bin sizes of 10°, and chains A (solid lines) and B (dashed lines) of OXA-48 are plotted separately. 6α-hydroxyethyl* *conformations I, II and III are labelled at 50°, 180°, and 290° respectively.*

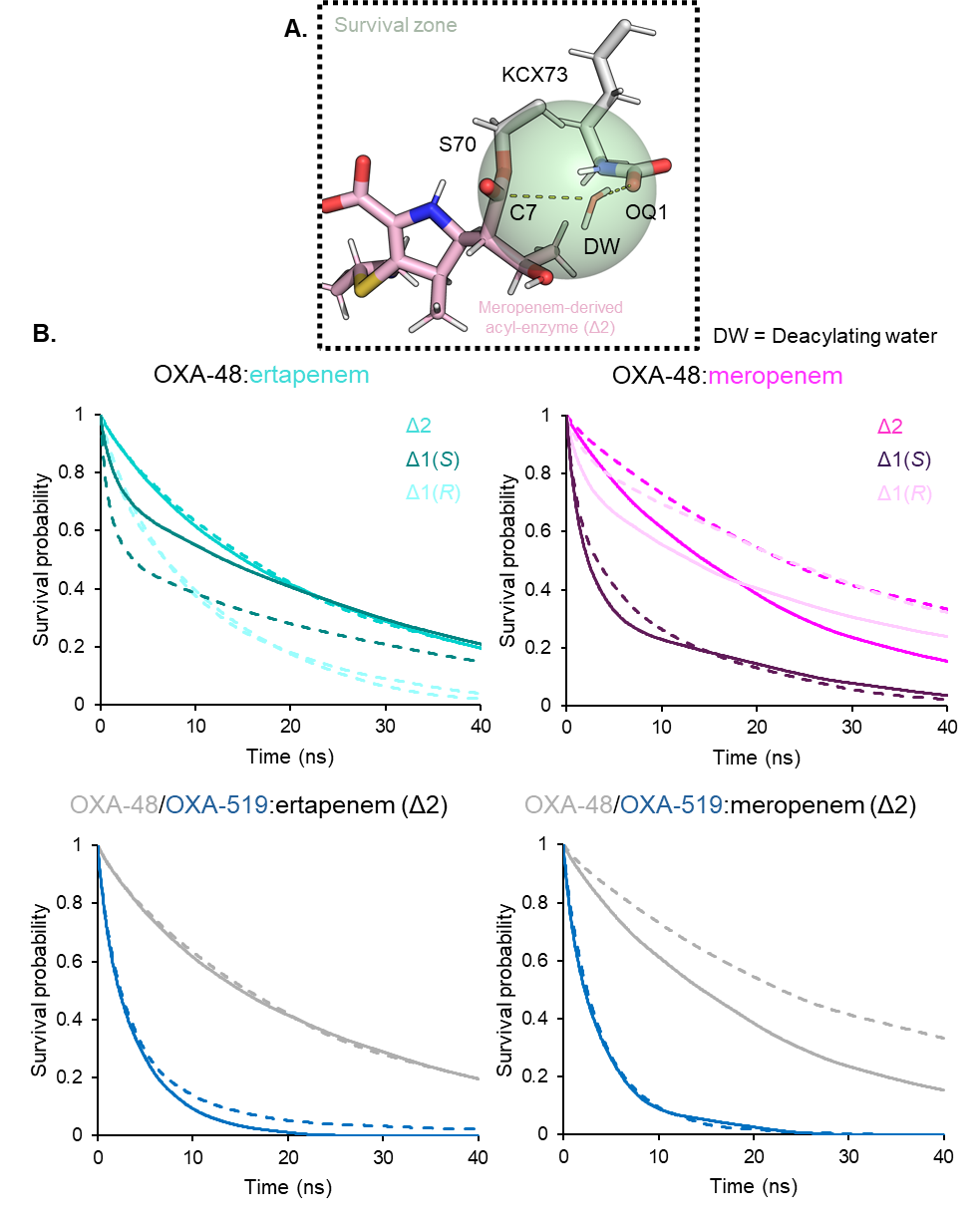
Figure S9: Survival probability of the deacylating water over simulations of meropenem and ertapenem-derived acyl-enzyme complexes with OXA-48 and OXA-519. *(A) Representative snapshot of OXA-48:meropenem (Δ2) from MD simulations displaying the survival zone (green sphere) for a water (DW) deemed to be in a deacylating position. The sphere has a radius of 3 Å between the C7 carbon of the acyl-enzyme complex and KCX73^OQ1^. (B) Survival times of waters within the survival zone of OXA-48 and OXA-519 acyl-enzyme complexes with meropenem and ertapenem. Solid lines are for chain A complexes and dashed lines for chain B.*

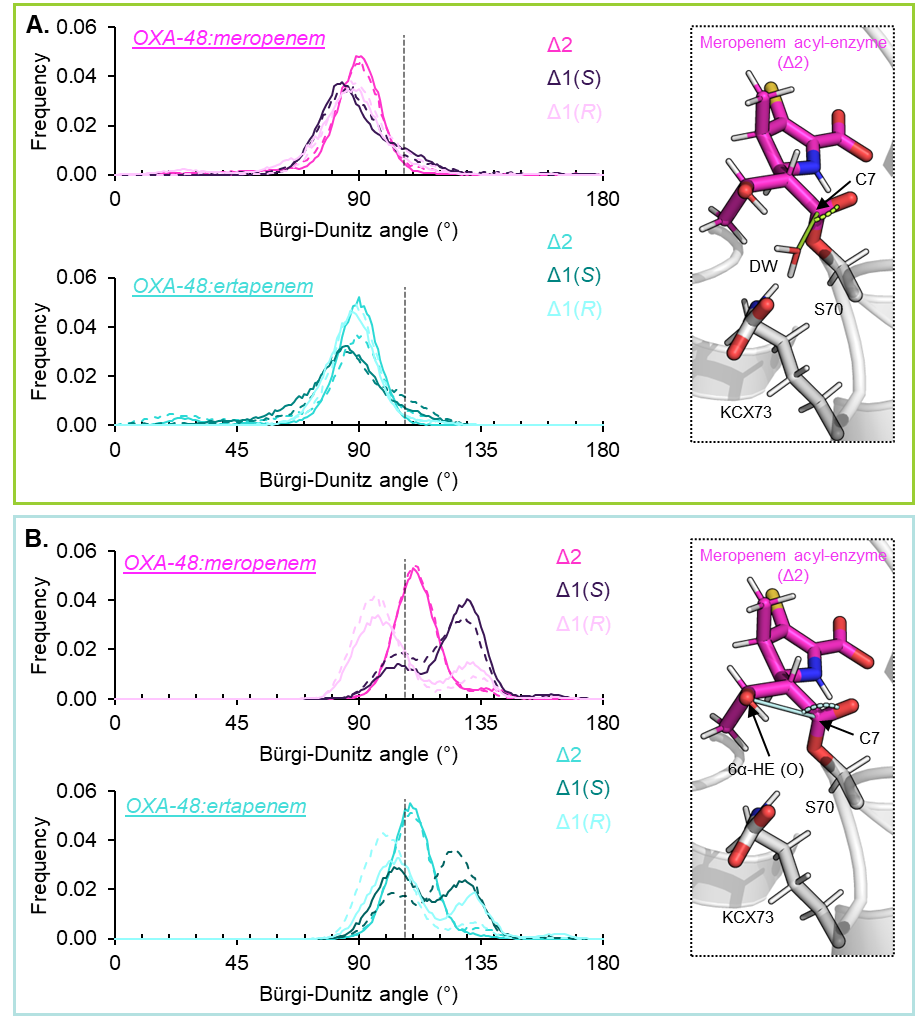
Figure S10: Bürgi-Dunitz angle for hydrolysis and β-lactone nucleophilic attack trajectories over MD simulations of meropenem- and ertapenem-derived acyl-enzyme complexes with OXA-48. *Frequency histograms of the Bürgi-Dunitz angle for (A.) hydrolysis and (B.) β-lactone deacylation during MD simulations of OXA-48:carbapenem acyl-enzymes. The measured angles are shown in the adjacent panel, on a snapshot taken from OXA-48:meropenem (Δ2) MD simulations, with the acyl-enzyme depicted as purple sticks. The deacylating water (DW), carbapenem 6α-hydroxyethyl oxygen (6α-HE (O)) and C7 carbon are labelled. Angles were separated into bin sizes of 1°, and chains A (solid lines) and B (dashed lines) of OXA-48 are plotted separately. The optimal Bürgi-Dunitz angle (107°)*^6^ *is overlayed as a grey dashed line on each frequency histogram.*

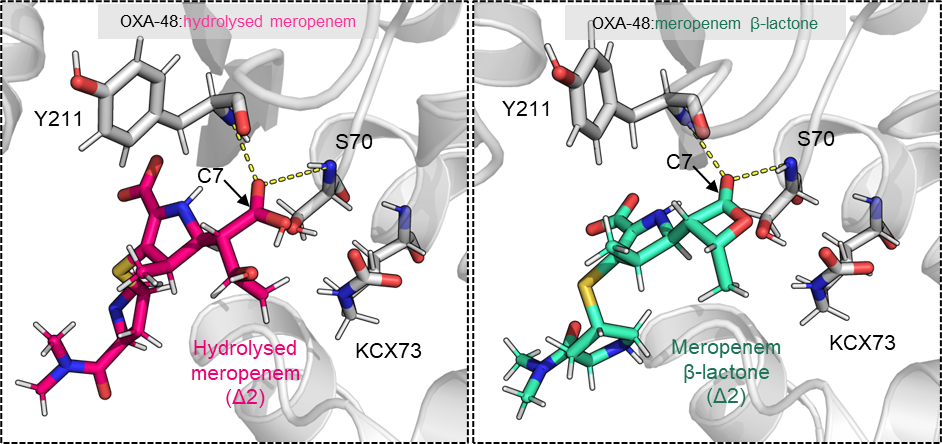

Figure S11: Starting conformations of hydrolysed and β-lactone meropenem products for MD simulations. *Snapshots were taken after heating and pressure equilibrations and prior to 500 ns productions runs (see Hoff* et al.^4^ *for energy minimisation and equilibration step details). The oxyanion hole is detailed by backbone amide nitrogens of Ser70 and Tyr211.*

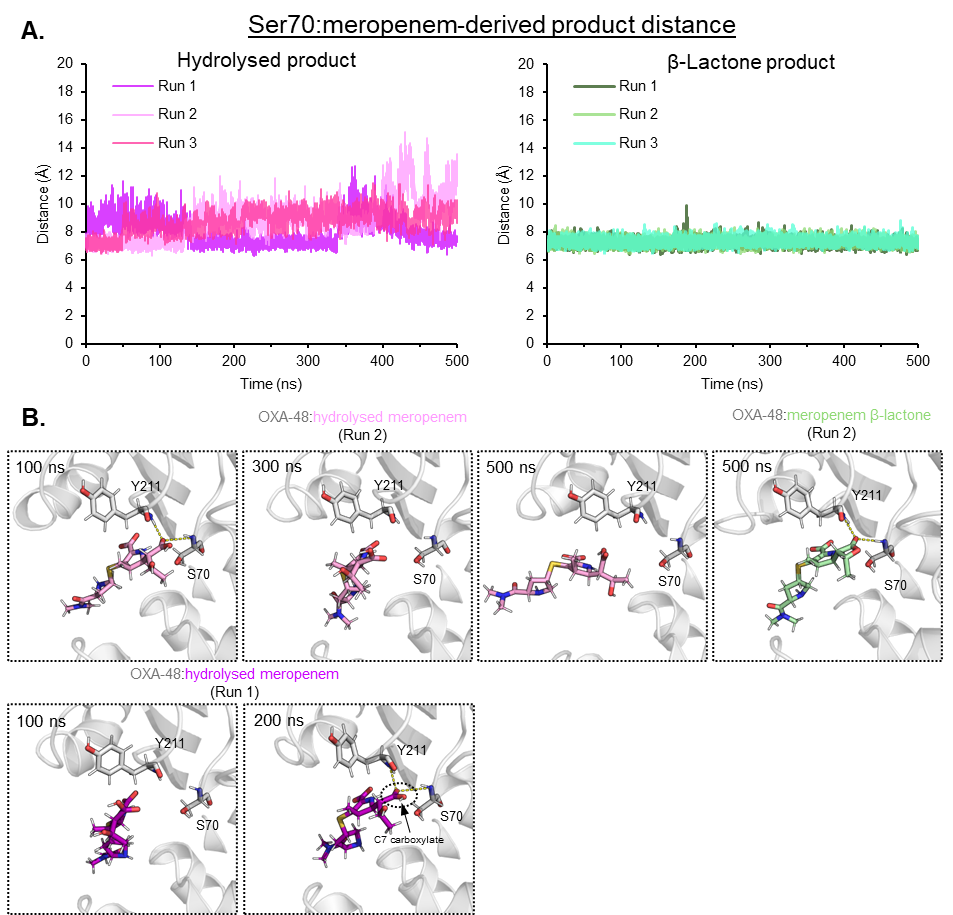

Figure S12: MD simulations of OXA-48 in complex with nascent meropenem-derived hydrolysed and β-lactone products. *(A) Ser70:meropenem product distances (centre of mass) over each 500 ns simulation trajectory. (B) Simulations snapshots at different time-points corresponding to repeats from panel A. OXA-48 is shown in grey and meropenem-derived hydrolysed and β-lactone products as pink/purple and green, respectively.*

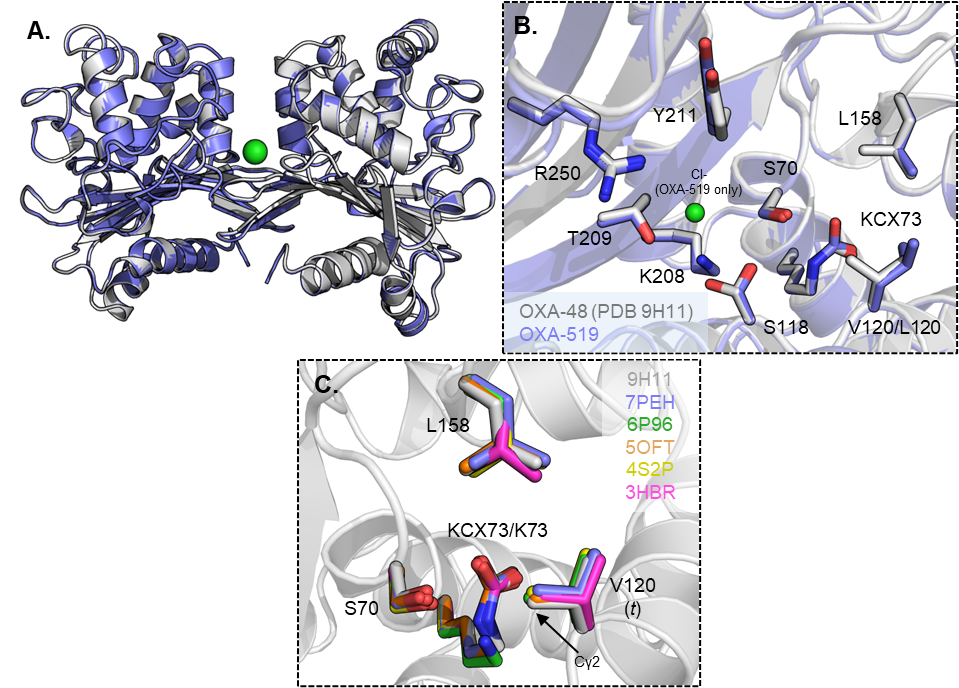

Figure S13: Overlays of uncomplexed OXA-48 and OXA-519. *(A) Overlay of uncomplexed OXA-48 (grey) (PDB 9H11) and OXA-519 (blue), solved in the same space group (*P*6_5_22) and under similar conditions*^4^*. (B) Chain A active site overlays with side-chains of some important residues shown as sticks. Chloride ions are indicated by green spheres. (C) Overlay of the deacylating-water channels of PDB-deposited uncomplexed OXA-48 structures (PDB 9H11*^4^*, 7PEH*^5^*, 6P96*^7^*, 5OFT*^8^*, 4S2P*^9^*, 3HBR*^10^*), showing only chain A active sites. The χ_1_ rotamer form of Val120 is in parentheses.*

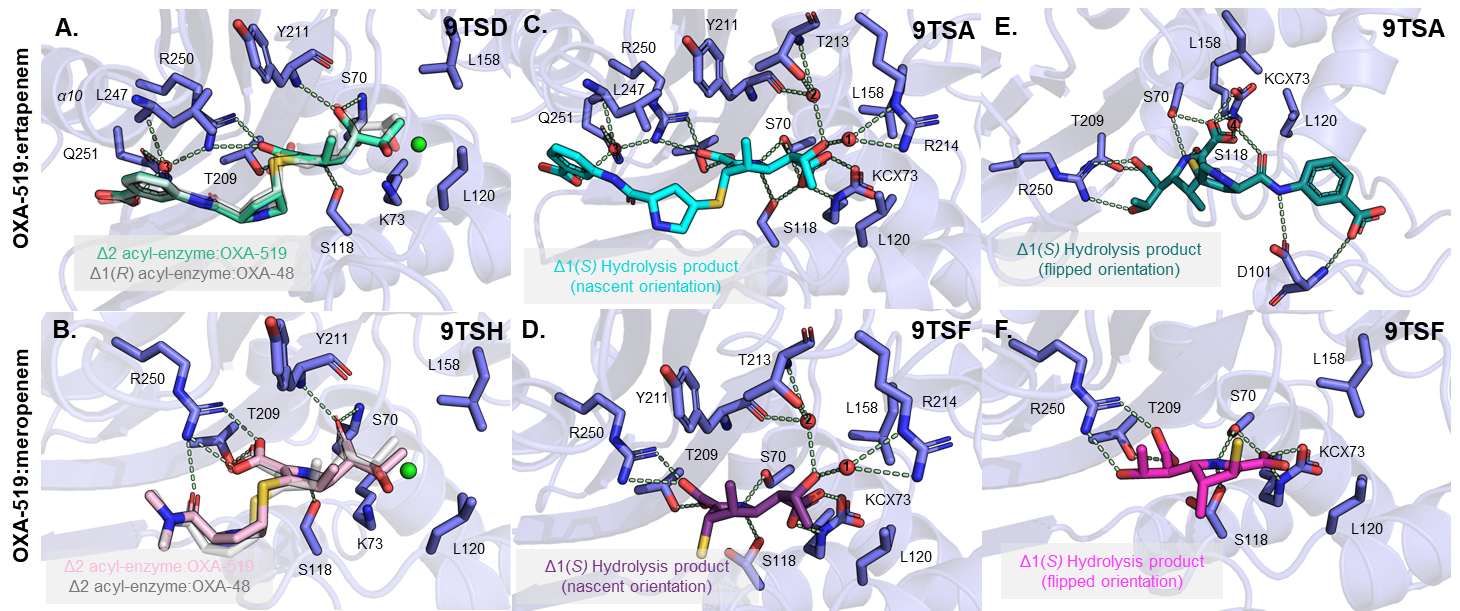
Figure S14: Comparison of hydrogen bond networks in the active site of the OXA-519 meropenem-derived acyl-enzyme complexes and hydrolysis products. *OXA-519:ligand interactions within hydrogen bonding distance are shown as pale green dashes. Bridging waters are shown as red spheres and active site chloride ions as green spheres. (A) OXA-519:ertapenem (22 hours, chain A), (B) OXA-519:meropenem (22 hours, chain A), (C) OXA-519:ertapenem (30 minutes, chain A, nascent orientation), (D) OXA-519:meropenem (2 hours, chain A), (E) OXA-519:ertapenem (30 minutes, chain B), (F) OXA-519:meropenem (2 hours, chain B). OXA-48:ertapenem (1 hour soak, chain A, Δ1(R) tautomer, PDB 9TSK) and OXA-48:meropenem (1 hour soak, chain A, Δ2 tautomer, PDB 9TSI) are overlayed as transparent grey sticks with their respective complexes in OXA-519 (****Fig. S2****).*

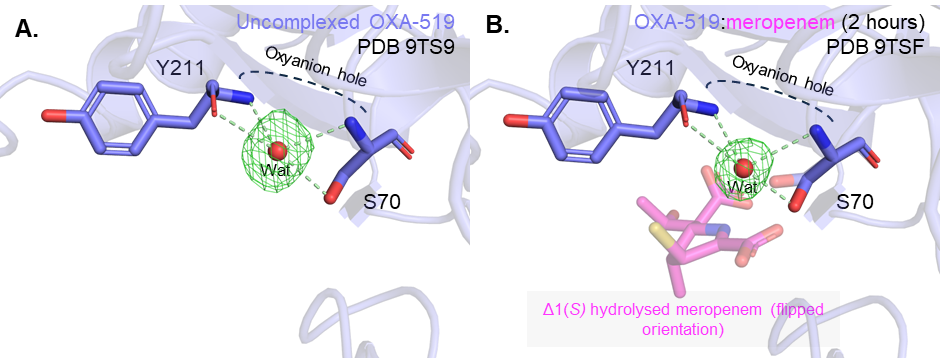

Figure S15: Unliganded and hydrolysed carbapenem-bound OXA-519 oxyanion hole:water interactions. *Chain B oxyanion hole of (A) OXA-519 apoenzyme and OXA-519:meropenem (2 hours) (both chain B). Water* F*_o_-*F*_c_ omit map (contoured to 3σ) is shown as green mesh, and hydrogen bonding interactions with OXA-519 shown as green dashed lines. Dual-occupancy molecules are shown as transparent sticks.*

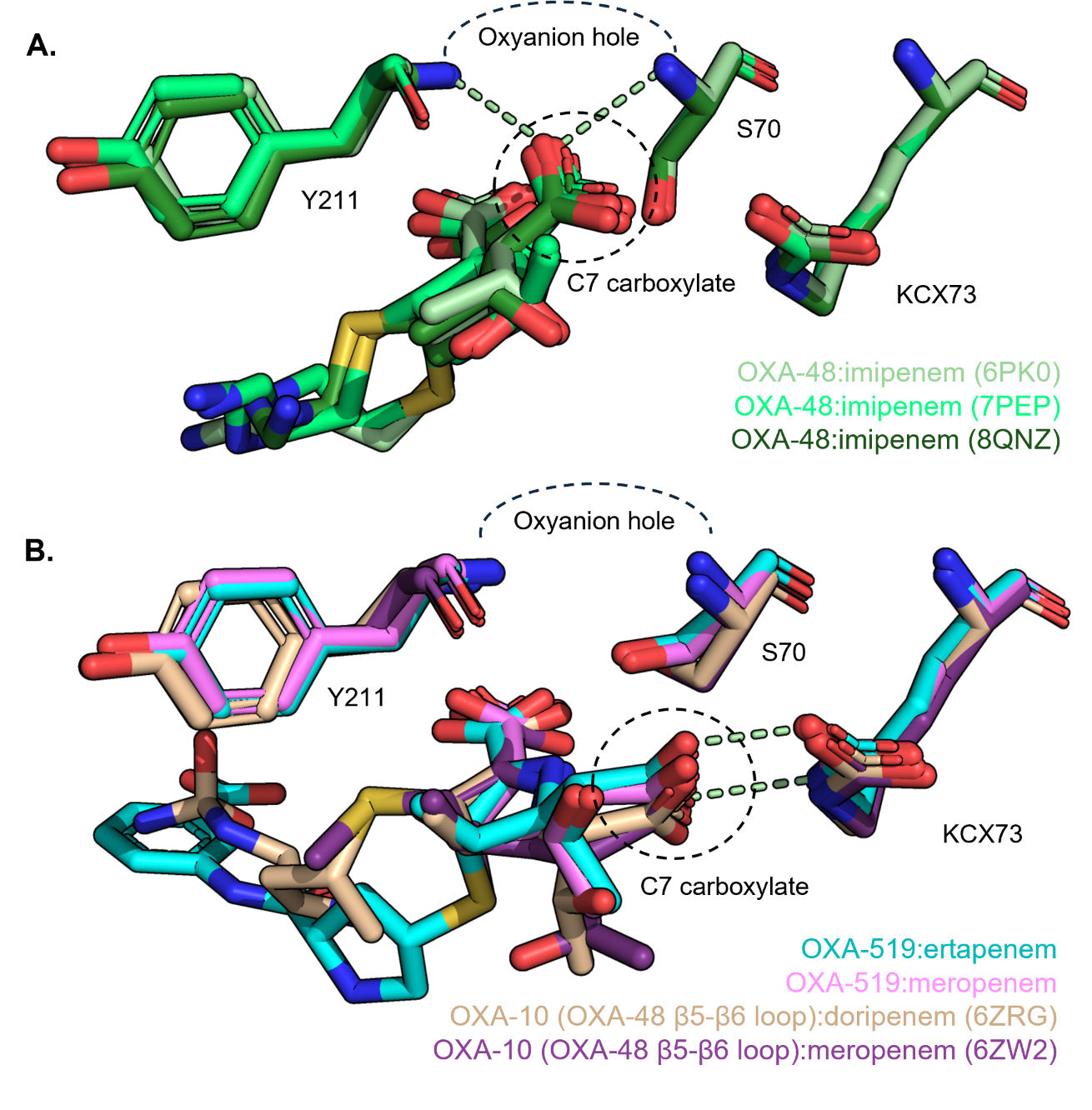
Figure S16: Overlay of structures of class D SBLs in complex with carbapenem and penem hydrolysed products in the initially observed orientation. *(A) Hydrolysed carbapenem structures where the C7 carboxylate is positioned in the oxyanion hole created by Tyr211 and Ser70: OXA-48:imipenem (6PK0*^11^*), OXA-48:imipenem (7PEP*^5^*), OXA-48:imipenem (8QNZ*^5^*). (B) Hydrolysed carbapenem/penem structures in the initially observed orientation, where the C7 carboxylate is positioned outside the oxyanion hole towards Lys73: OXA-10 (substituted OXA-48 β5 - β6 loop*^12^*):doripenem (6ZRG), OXA-10 (substituted OXA-48 β5 - β6 loop):meropenem (6ZW2), OXA-519:ertapenem (this work, 30 minute soak), OXA-519:meropenem (this work, 2 hour soak). Possible hydrogen bonding interactions between the C7 carboxylate and active site of OXA-519 are indicated by green dashed lines.*

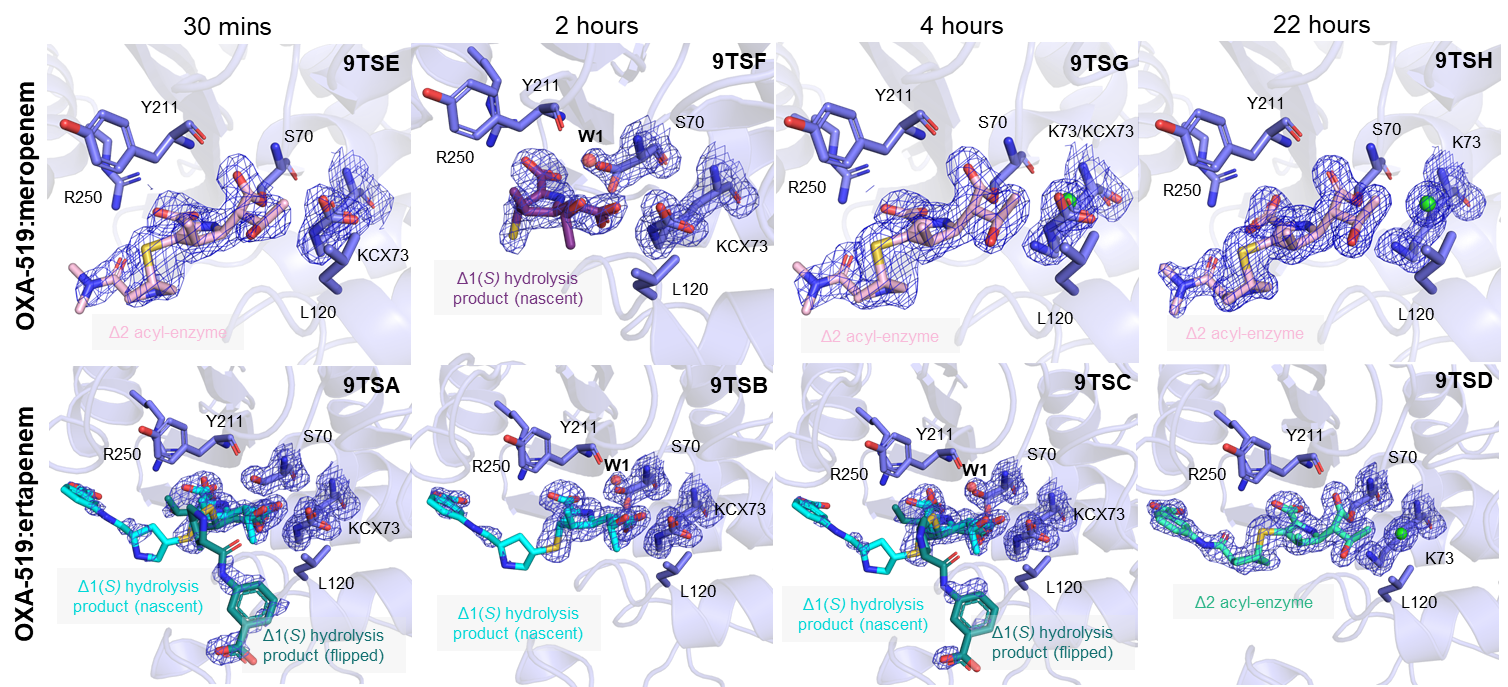

Figure S17: Time-course crystallography of meropenem and ertapenem complexes with OXA-519. *Chain A active sites of crystal structures of OXA-519 at carbapenem soaking time points of 30 minutes, 2 hours, 4 hours and 22 hours. Selected active site residues are shown as blue sticks and meropenem- and ertapenem-derived products as sticks in shades of pink and turquoise respectively. Chloride ions are represented as green spheres, waters as red spheres. Final model 2*F*_o_-*F*_c_ electron density maps of ligands, Ser70, Lys73 and active site chloride ions are shown as a blue mesh, contoured to 1σ.*

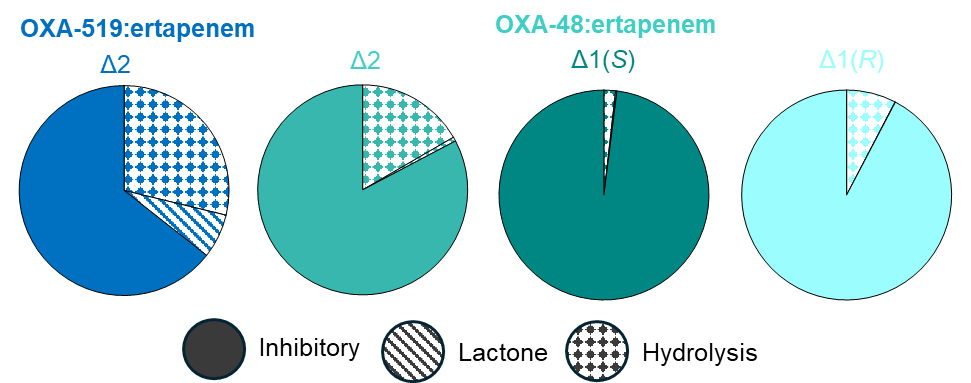

Figure S18: Conformational sampling of ertapenem-derived acyl-enzymes over MD simulations of OXA-48 and OXA-519. *Proportion of OXA-48/OXA-519:ertapenem acyl-enzyme simulations in either the β-lactone forming or hydrolysis-promoting conformations, averaged over both active sites of each enzyme. The geometric parameters that define these conformations are described in* ***Fig. 6*** *and displayed in* ***Figs. S19 and S20****.*

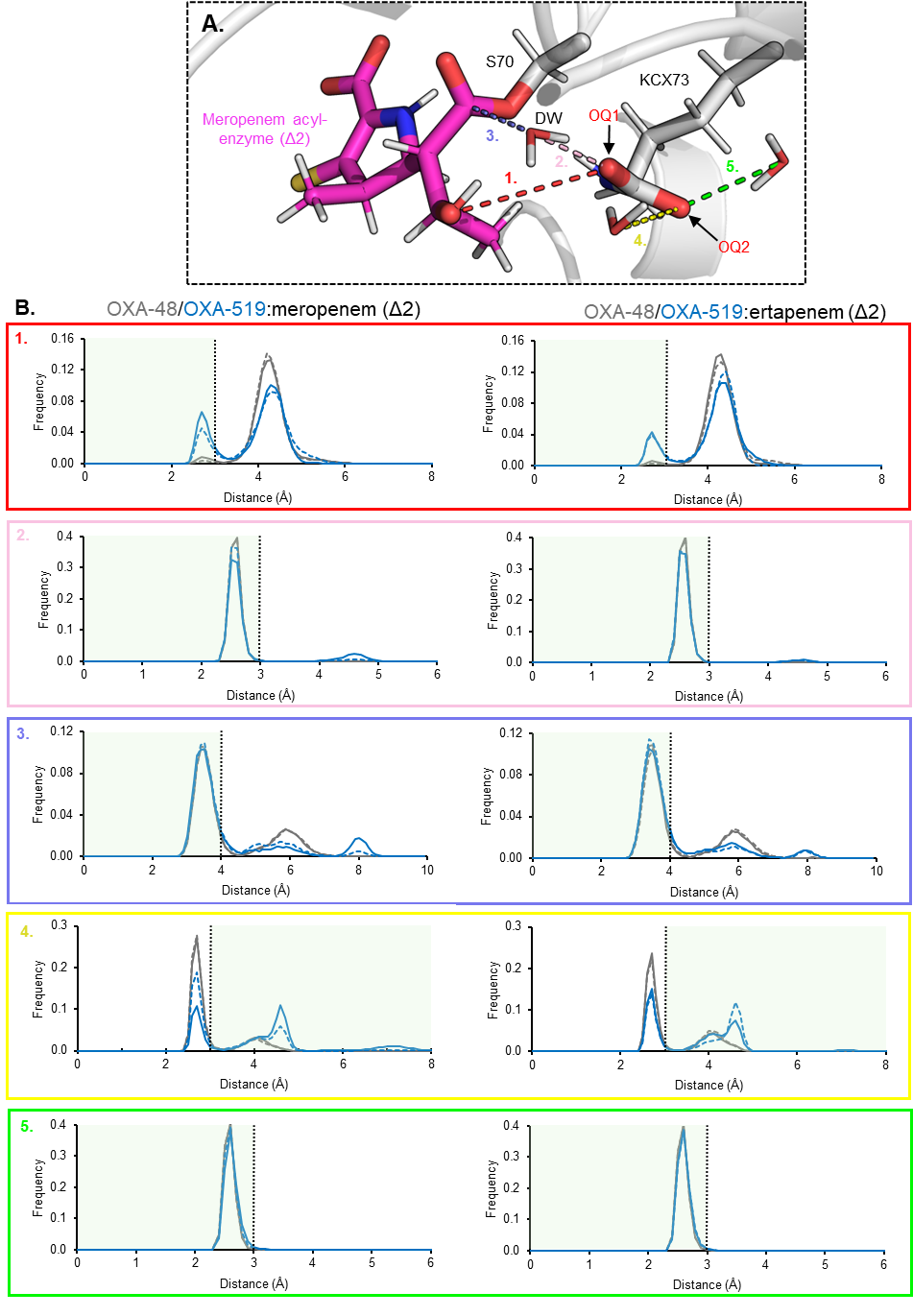
Figure S19: Carbapenem acyl-enzyme complex distance parameters used to filter MD simulations for deacylation promoting conformations. *(A) A simulation snapshot with each of the distance parameters (1 to 5) chosen to filter MD trajectories for hydrolysis- (parameters 2 to 5) and β-lactone (parameters 1, 4 and 5) promoting acyl-enzyme conformations. Angle parameters shown in* ***Fig. S20*** *were also included in the filtering analysis. (B) Frequency histograms of each of these distance parameters for simulations of meropenem and ertapenem acyl-enzymes (Δ2-enamine) with OXA-48 (blue) and OXA-519 (grey). Full lines are for chain A complexes, dashed lines for chain B complexes. Areas shaded in green represent distances that would be favourable for deacylation by hydrolysis or β-lactone formation.*

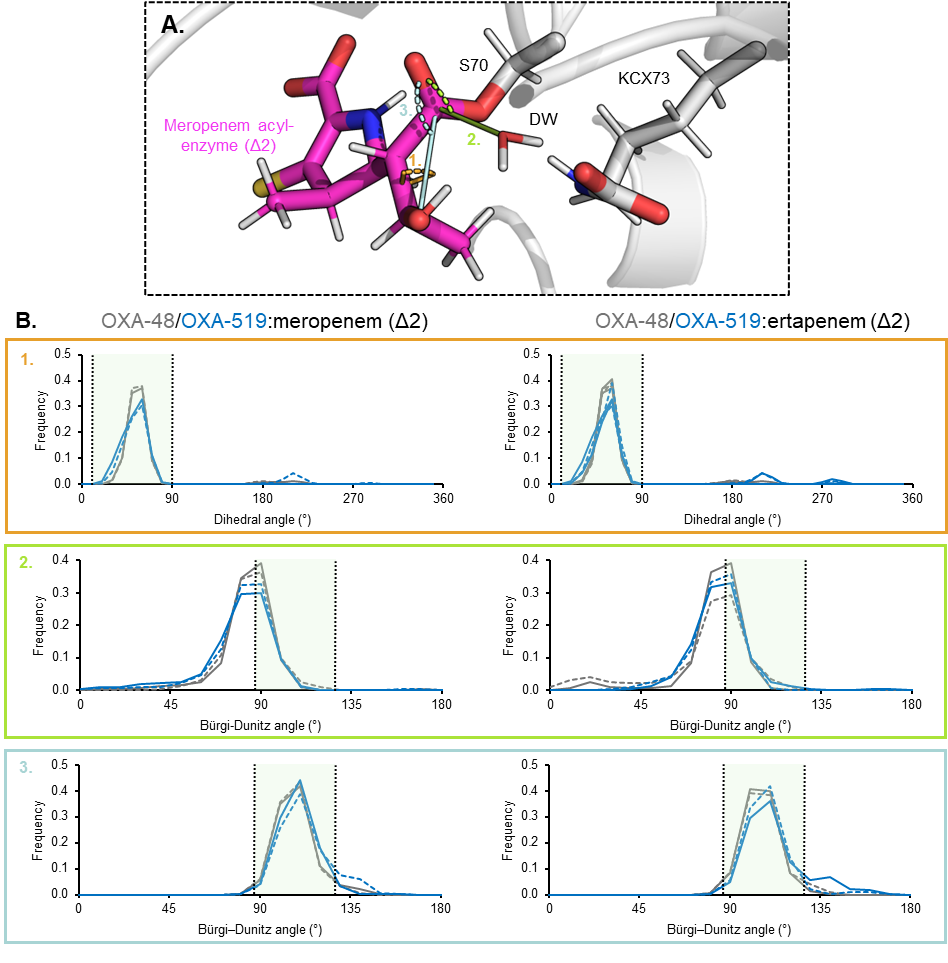

Figure S20: Carbapenem acyl-enzyme complex angle parameters used to filter MD simulations for deacylation promoting conformations. *(A) A simulation snapshot with each of the dihedral (parameter 1, C7-C6-C8-O 6α-hydroxyethyl angle) and Burgi-Dünitz angle parameters (2 and 3) chosen to filter MD trajectories for deacylation promoting conformations. Parameters 1 and 2 were chosen for hydrolysis-promoting acyl-enzyme conformations and parameters 1 and 3 for β-lactone-promoting conformations. Distance parameters shown in* ***Fig. S19*** *were also included in the filtering analysis. Parameter (B) Frequency histograms of each of these angle parameters for simulations of meropenem and ertapenem acyl-enzymes (Δ2-enamine) with OXA-48 (blue) and OXA-519 (grey). Full lines are for chain A complexes, dashed lines for chain B complexes. Areas shaded in green represent angles that would be favourable for deacylation by hydrolysis or β-lactone formation, with the preferred Bürgi-Dunitz angle for nuclophilic attack being 107±20°* ^6^*.*

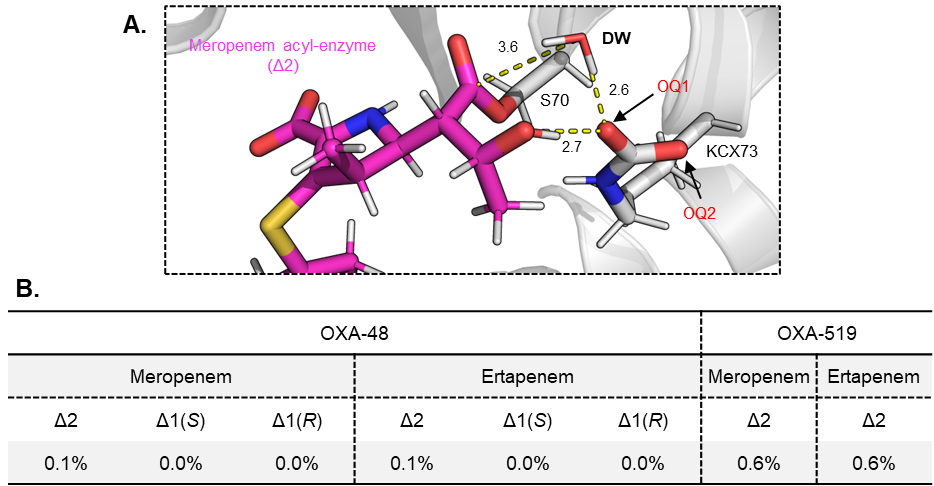

Figure S21: Observation of carbapenem-derived acyl-enzymes in “mixed” hydrolysis and β-lactone promoting conformations. *(A) Representative snapshot of the OXA-48:meropenem acyl-enzyme complex (in the Δ2-enamine configuration, pink) in a conformation that satisfies all the distance and angle criteria from* ***Figs. S19 and S20****. This is indicated by the 6α-hydroxethyl oxygen proximal to KCX73 (OQ1) and a water (DW) in a deacylating position adjacent to the carbapenem C7 carbon. (B) Proportion of OXA-48 and OXA-519 simulations as carbapenem-derived acyl-enzymes in this “mixed” conformation, averaged across both active site complexes.*

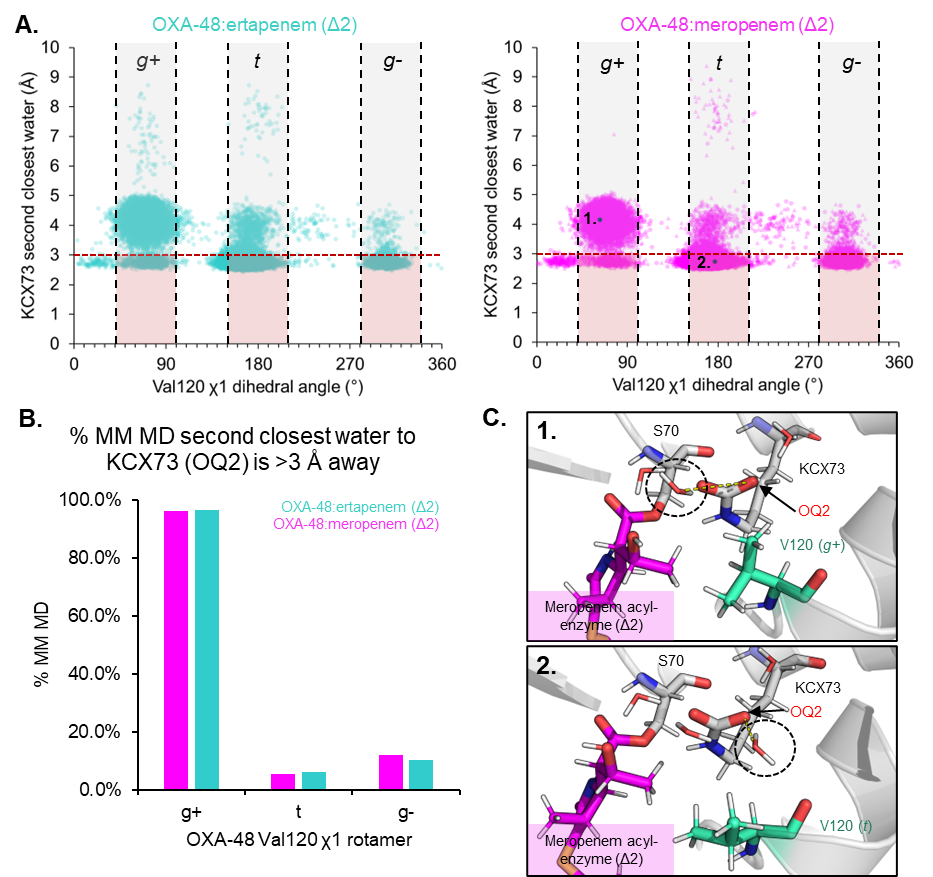
Figure S22: Relationship between Val120 rotamer dynamics and water associated with carboxylated Lys73. *(A) Scatter plots of OXA-48:meropenem/ertapenem derived acyl-enzyme complex (Δ2) MD simulations with Val120 χ_1_ dihedral (N-Cα-Cβ-Cγ1) angles (°) against the distance between KCX73 (OQ2) and a second closest water (O) molecule. Each dot represents a snapshot taken at 40 ps intervals during the simulations, with circles for chain A complexes and triangles for chain B complexes. Val120 χ_1_ rotamer forms are highlighted in between the black dashed lines, with the red dashed lines representing an arbritarily chosen distance cutoff for the second closest water to be sufficiently close to KCX73 (OQ2) to hydrogen bond and therefore impede deacylation*^13^*. (B) Proportion of OXA-48:carbapenem acyl-enzyme simulations where the second closest water to KCX73 (OQ2) is outside hydrogen bonding distance (3 Å). (C) Snapshots of OXA-48:meropenem acyl-enzyme (Δ2) simulations showing Val120 in (1.)* g+ *and (2.)* t *χ_1_ rotamers. The second closest water distance to KCX73 (OQ2) is indicated by a yellow dashed line.*

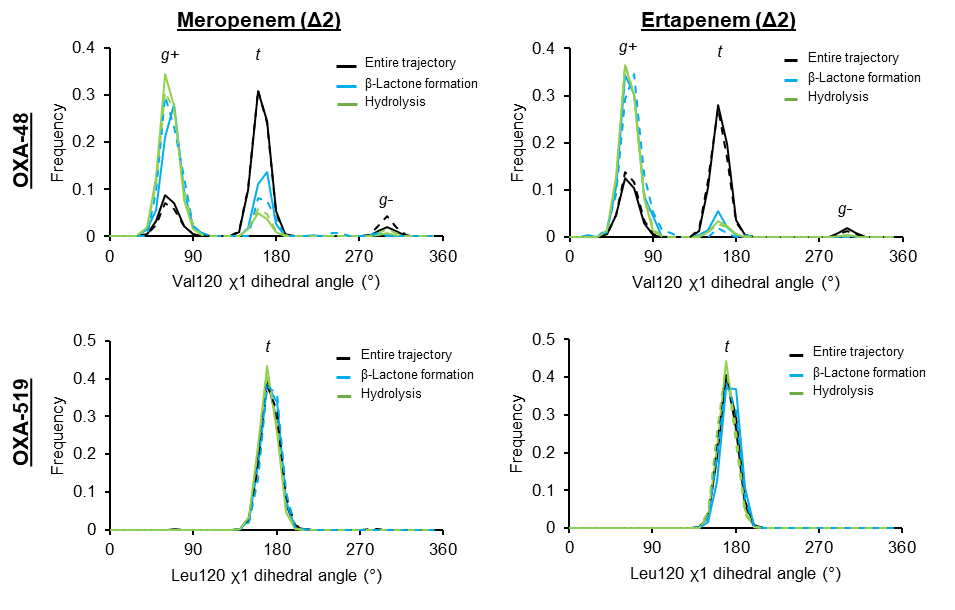
Figure S23: Position 120 side-chain rotamer conformational sampling during MD simulations of OXA-48 and OXA-519 meropenem and ertapenem-derived acyl-enzymes. *Frequency histograms of Val120 (OXA-48) and Leu120 (OXA-519) χ_1_ dihedral (N-Cα-Cβ-Cγ1) angles (°) during MD simulations of ertapenem- and meropenem-derived acyl-enzymes in the Δ2-enamine tautomer forms. Black lines represent residue-120 χ_1_ dihedral sampling over the entire simulation trajectories, whereas blue and green lines are from simulation frames where the acyl-enzyme complex is in either β-lactone (blue) or hydrolysis (green) promoting deacylating conformation. These conformations were filtered using the parameters described in* ***Figs. S19 and S20****. Full lines are for chain A complexes, dashed lines are for chain B complexes.*

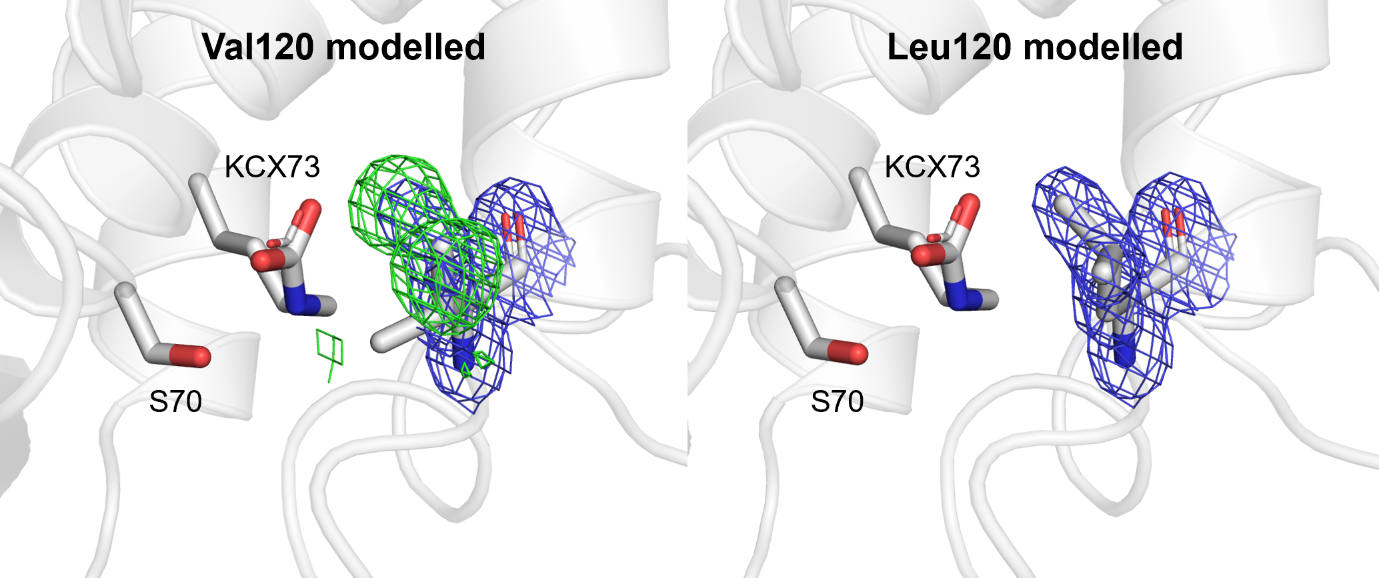

Figure S24: Modelling of Leu120 in OXA-519. *Active site views of the OXA-519 apo-enzyme where (A) Val120 was modelled following initial rigid body refinement using unliganded OXA-48, and where (B) Leu120 was modelled instead, as in the final refined structure of OXA-519. Residue 120* F­*_o_-*F*_c_ (green mesh, contoured to 3σ) and 2*F­*_o_-*F*_c_ (blue mesh, contoured to 1.5σ) maps are shown.*

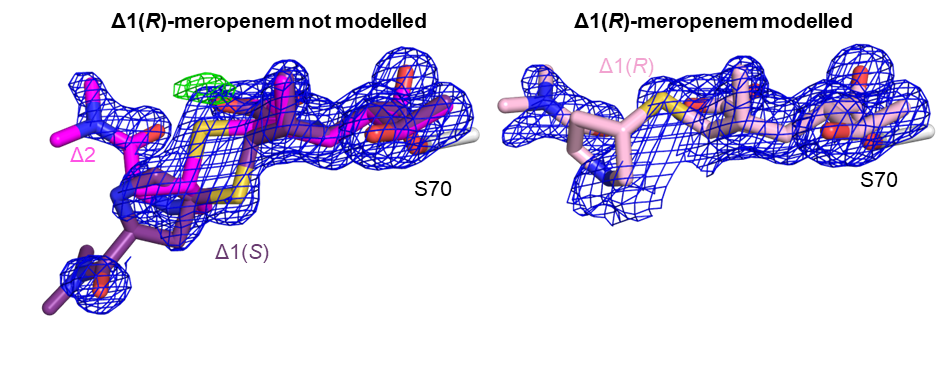
Figure S25: Attempted modelling of the meropenem Δ1(*R*)-tautomer into the chain A active site of OXA-48 at 2 hours. *Ser70 (grey), Δ2-meropenem (magenta), Δ1(*S*)-meropenem (purple), Δ1(*R*)-meropenem (pink) are shown as sticks. The combined 2*F*_o_-*F*_c_ map of meropenem complexes is shown as a blue mesh (contoured to 1σ) and the* F*_o_-*F*_c_ difference map (contoured to 3σ and carved to 2 Å around the C2-sulphur of Δ2 meropenem) is shown as a green mesh.*
